# Depletion of CX3CR1^+^ macrophages results in disrupted functionality and immune surveillance within epididymis and testis

**DOI:** 10.1101/2025.09.26.678714

**Authors:** D Ai, L Kreyling, MA Battistone, ML Elizagaray, A Chen, S Bhushan, M Fijak, M Speckmann, G Michel, T Procida-Kowalski, M Bartkuhn, M Sprang, JU Mayer, A Meinhardt, C Pleuger

**Affiliations:** Department of Anatomy and Cell Biology, Unit of Reproductive Biology, Justus-Liebig University Giessen, Germany; Hessian Centre of Reproductive Medicine, Justus-Liebig University Giessen, Germany; Program of Membrane Biology, Nephrology Division, Department of Medicine, Massachusetts General Hospital and Harvard Medical School, Boston, MA, USA; Flow Cytometry Core Facility, Justus-Liebig University Giessen, Germany; Biomedical Informatics and System Medicine, Science Unit for Basic and Clinical Medicine, Justus-Liebig University Giessen, Germany; Department of Dermatology, University Medical Center of the Johannes Gutenberg-University Mainz, Mainz, Germany; Research Center for Immunotherapy (FZI), University Medical Center of the Johannes Gutenberg-University Mainz, Mainz, Germany; Institute of Quantitative & Computational Biosciences, Johannes Gutenberg University Mainz, Mainz, Germany

**Keywords:** Male reproductive tract, epididymis, testis, CX3CR1^+^ macrophages, mononuclear phagocytes, mucosal immunology, immune regulation, sperm function, spermatogenesis, epithelial physiology, epididymitis, UPEC

## Abstract

A finely tuned immune regulation within the epididymis and testis is essential for male reproductive health. This balance is especially critical in the epididymis, where sperm mature and ascending infections frequently disrupt homeostasis, resulting in regionally different immune responses and potential long-term fertility impairments. We previously demonstrated that the epididymis harbors a region-specific immunological scaffold, with CX3CR1^+^ macrophages as the most prominent epithelium-associated immune cell population. Here, we established a transgenic mouse model to selectively deplete these intraepithelial CX3CR1^+^ macrophages within the epididymis, resulting in focal epithelial damage and impaired sperm maturation processes essential for proper sperm functionality. Additionally, a mild reduction of the testicular macrophage pool resulted in transient disruptions in spermatogenesis and steroidogenesis. Although the macrophage niche was repopulated after depletion, the newly recruited cells displayed altered phenotypes consistent with persistent sperm alterations. Following infection with uropathogenic *Escherichia coli* (UPEC), macrophage-depleted mice exhibited exacerbated immune responses - particularly in normally protected proximal epididymal regions - with earlier onset and more severe tissue damage. Transcriptomic analysis revealed a failure to restrain inflammatory responses, especially in genes involved in immune regulation and antibacterial defense, accompanied by elevated immune cell infiltration in infected macrophage-depleted mice. Overall, our findings confirm a crucial role for CX3CR1⁺ macrophages in preserving epithelial integrity and modulating immune responses, supporting a stable tissue environment necessary for efficient organ function of both epididymis and testis.

**Significance statement:** Maintaining immune balance in the epididymis is essential for tissue health and protection against ascending infections. Using a transgenic mouse model that allows for selective depletion of CX3CR1⁺ macrophages, this study examines their role in both the epididymis and testis under normal and infectious conditions. The results show that the removal of these macrophages causes localized epithelial damage, changes in immune cell make-up, and increased inflammation in the epididymis after bacterial infection, while also causing mild, reversible problems with spermatogenesis and steroid production in the testis. These findings support the idea that CX3CR1⁺ macrophages contribute to region-specific immune regulation and epithelial stability—key features for keeping the tissue environment suitable for proper sperm development.

## Introduction

Male fertility relies on the coordinated function of the testis and epididymis—adjacent organs with distinct roles in sperm production and maturation. Although both organs are exposed to similar antigens and immunological challenges, they exhibit markedly different immune architectures. Spermatogenesis occurs within the immune-privileged environment of the testis, whereas spermatozoa, once released, encounter the immune system in the epididymis, where essential post-testicular maturation takes place^1^. To prevent autoimmune reactions against autoantigenic spermatozoa the immune system must establish a tolerogenic milieu within the epididymis while retaining the capacity to mount effective immune responses against ascending urogenital pathogens.

Disruption of this delicate immunological balance – most often triggered by acute bacterial infections^2^ - often results in long-term fertility impairments. Indeed, up to 40% of patients with epididymitis experience persistent sub- or infertility^3^, with a clear correlation between epididymitis and the occurrence of anti-sperm antibodies^4^. Experimental mouse models have shown that epididymal tissue damage is primarily driven by an exacerbated proinflammatory immune response rather than bacterial presence *per se*^5–7^. Notably, these immune responses manifest most severely in the cauda epididymidis, leading to duct stenosis/obstruction and fibrotic remodeling^8,9^, whereas in proximal regions inflammation is thoroughly controlled and rapidly resolved, preserving tissue integrity^7^.

Recent studies have highlighted the epididymis as a mucosal tissue with a regionally specialized immune architecture. Resident immune cells, including both myeloid and lymphoid populations, are strategically positioned along the duct^7,10,11^, occupying specific niches that reflect regional demands for tissue homeostasis, sperm tolerance and immune defense. Among these, CX3CR1^+^ mononuclear phagocytes, with a macrophage signature, represent the predominant immune cell population and are primarily located within and around the epididymal epithelium^7,11–13^. Particularly within the initial segment, the entry point for sperm, intraepithelial CX3CR1^+^ macrophages extend long protrusions between neighboring epithelial cells toward the lumen, suggesting a crucial sentinel role in monitoring luminal contents^11,12^

Despite their strategic positioning and unique morphology, the functional role of CX3CR1^+^ macrophages in maintaining epididymal immune balance remains incompletely understood. Transcriptomic profiling reveals that epididymal CX3CR1^hi^ macrophages typically display a homeostatic transcriptional signature, in contrast to the more proinflammatory profile of interstitial macrophage subsets^7^. Across different epididymal regions, CX3CR1^hi^ macrophages become less prevalent and adopt a more pro-inflammatory signature towards the distal end^11^, consistent with the regiońs heightened vulnerability to pathogen invasion and tissue damage. These observations point to a dual role for CX3CR1^+^ macrophages within the epididymis – supporting epithelial integrity and sperm maturation within proximal regions, while serving as immune sentinels in distal regions. However, whether these cells are causally required for maintaining regional immune homeostasis and the consequences from their absence, have not been systematically examined.

Within the testis, macrophages substantially contribute to the establishment and maintenance of the immune-privileged environment that allows germ cells to develop without eliciting immune responses^14,15^. By actively suppressing and resolving inflammation (e.g. in response to ascending bacterial infections), they help preventing immune-mediated damage to developing gametes. Although testicular macrophages also (partially) express CX3CR1, they are functionally and transcriptionally distinct^6^ and are restricted to the interstitial space, in contrast to the more epithelium-associated localization observed in the epididymis.

In this study, we selectively depleted CX3CR1^+^ macrophages *in vivo* in a transgenic mouse model to investigate their role in regulating immune homeostasis in the epididymis and testis under physiological conditions. Given that macrophages in both the epididymis and testis express CX3CR1 and have been implicated in supporting sperm development and maturation^16,17^, we evaluated the impact of CX3CR1^+^ macrophage depletion on the local immune system and fertility-related physiological parameters in both organs. To further examine the functional consequences of macrophage loss during infection, we employed an established mouse model of acute bacterial epididymitis that closely mimics the clinical features observed in patients^18^ and analyzed region-specific immunity following infection with uropathogenic *Escherichia coli* (UPEC).

## Results

### Elaborating a mouse model to deplete intraepithelial CX3CR1^+^ macrophages within the epididymis

To investigate the role of intraepithelial macrophages in regulating epididymal immunity, we established a transgenic mouse model designed to predominantly target CX3CR1^hi^-expressing cells. This approach was based on previous findings showing that CX3CR1 is highly expressed by intraepithelial macrophages, with their abundance progressively declining from proximal to distal epididymal regions^7^.

Although interstitial CX3CR1^+^ macrophages and monocytes are also present in distal segments, these populations exhibit high turnover due to continuous recruitment from circulating monocytes^7^. To account for this dynamic exchange and ensure effective depletion of more stably resident intraepithelial subsets (excluding short-living neutrophils and monocytes), we implemented a two-week interval between tamoxifen (Tam)-induced diphtheria toxin receptor (DTR) expression and subsequent diphtheria toxin (Dtx) administration for targeted depletion in *Cx3cr1*^CreER^*Rosa26*^iDTR/tdTomato^ mice (**Fig. 1a**). To visualize and confirm successful Tam-induced recombination, a tdTomato (Tom) reporter allele was also incorporated into the Rosa26 locus. Following this regimen, we observed a significant reduction of Tom^+^ cells in both proximal and distal epididymal regions. A milder but still significant decrease was also detected in the testis (**Fig. 1b**). Across all examined tissues, F4/80^+^CX3CR1^+^ cells accounted for almost all Tom^+^ cells, confirming specific targeting of CX3CR1^+^ macrophages (**Fig 1c).** Administration of Tam or Dtx alone did not alter (i) other immune cell populations and (ii) the ratio of CX3CR1^+^F4/80^+^ cells within the CD45^+^ cells in either the epididymis or the testis (**Suppl. Fig. S1a**).

**Figure 1:**
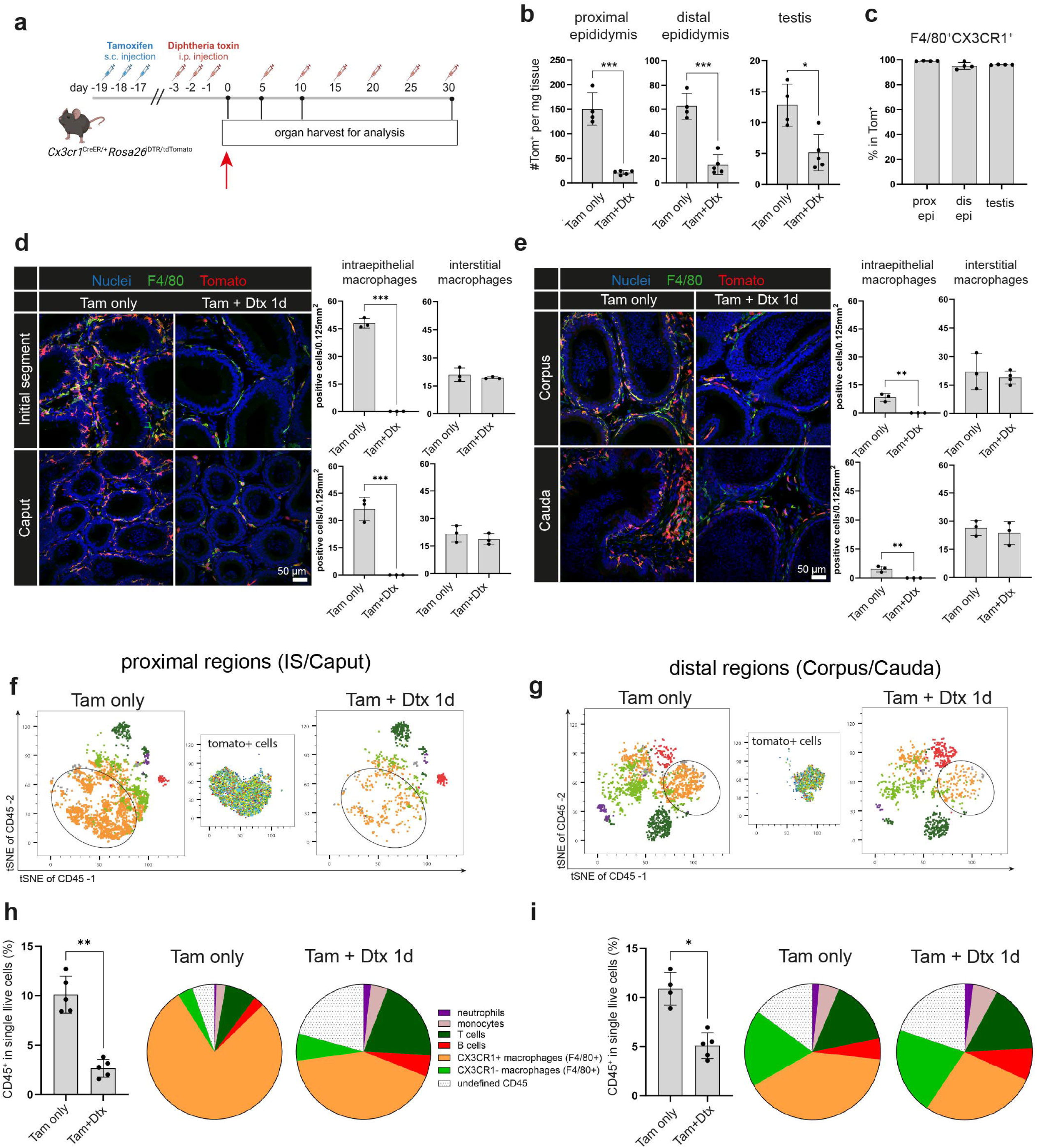
Targeted depletion of intraepithelial CX3CR1^+^ macrophages in the murine epididiymis. **(a)** Schematic overview of the depletion strategy using sequential tamoxifen (Tam) and diphtheria toxin (Dtx) treatment in *Cx3cr1*^CreER/+^*Rosa26*^iDTR/tdTomato^ mice. **(b)** Quantification of tdTomato^+^ (Tom^+^) cells per mg tissue in proximal (IS/ caput) and distal (corpus/ cauda) epididymis, and testis following Dtx-mediated depletion, assessed by flow cytometry one day after depletion. **(c)** Proportion of F4/80^+^CX3CR1^+^ cells among Tom^+^ cells (each data point represents pooled proximal and distal regions from left and right epididymis, n=4-5). **(d, e)** Confocal microscopy images of Tom^+^ cells (red) in intraepithelial and interstitial compartment of (D) proximal and (E) distal epididymal regions. Semi-quantitative analysis of Tom^+^ macrophages (F4/80^+,^ green) was performed by counting ≥5 representative areas per mouse (n=3). Nuclei were stained with DAPI (blue). **(f, g)** t-SNE plots (opt-SNE with iterations: 500, perplexity: 20) of CD45^+^ populations from representative Tam only and Tam+Dtx-treated mice, one day post depletion, in the (F) proximal and (G) distal epididymis. CD45^+^ cells were downsampled, gated as shown in Fig. S1E and overlaid onto the t-SNE plots. Tomato^+^ cells were gated separately and plotted on the t-SNE coordinates. Circles indicate differences in CX3CR1^+^ macrophages between Tam only and Tam+Dtx-treated mice. **(h, i)** Relative abundance of CD45^+^ cells and distribution of immune cell subsets (pie charts) within the CD45^+^ compartment in (H) proximal and (I) distal (G) regions of Tam only and Tam+Dtx-treated mice, one day post depletion. Immune cell populations are color-coded as in (F, G), based on the gating strategy in Fig. S1E. Data are presented as mean ± SD from two independent experiments (n=4-5 mice). Statistical analysis was performed using Mann-Whitney *U* test, *p<0.05, **p<0.005, ***p<0.001).

Confocal microscopy revealed that in proximal regions (initial segment, caput) Tom^+^ cells were predominantly intraepithelial/ epithelia-associated macrophages (**Fig. 1d**). In distal regions (corpus, cauda), Tom+ cells included both intraepithelial and a substantial number of interstitial macrophages (**Fig. 1e**). However, Dtx administration selectively reduced intraepithelial macrophages across all regions, while interstitial CX3CR1^+^ macrophages were only mildly affected (**Fig. 1d,e**), consistent with our strategy to preferentially deplete the more stable resident subsets.

High-dimensional analysis of CD45^+^ cells confirmed that tdTomato fluorescence was selectively restricted to CX3CR1^+^ macrophages, with no detectable expression in other investigated populations (including CX3CR1^-^ macrophages, monocytes, neutrophils, T cells or B cells) confirming the specific targeting of macrophages. This Tom^+^CX3CR1^+^ macrophage population was strongly diminished following Dtx administration (**Fig. 1f,g, Suppl. Fig. S1b**). No such changes occurred with Tam or Dtx treatment alone (**Suppl. Fig. S1c**).

Overall, the significant decrease in CD45^+^ cell numbers observed in both proximal and distal regions was primarily attributable to the loss of CX3CR1^+^ macrophages, while other immune cell populations remained quantitatively stable (**Fig. 1h,i**). Notably, the macrophage depletion was accompanied by a relative increase in neutrophils and T cells, suggesting a compensatory shift of pro-inflammatory immune cells in the local immune landscape, particularly within proximal regions of the epididymis (**Suppl. Fig S1d**).

### Loss of CX3CR1^+^ macrophages results in focal epithelial damage and impaired sperm maturation within the epididymis

Although overall epididymal morphology appeared unaffected immediately after Dtx administration (**Suppl. Fig. S2a**), CX3CR1^+^ macrophage-depleted mice developed pronounced focal tissue damage peaking at 10 days after Dtx-mediated depletion, particularly in segments S2, S5/6 and S8 (**Fig. 2a**). Surprisingly, no overt epithelial damage was observed in the initial segment (**Fig. 2a**), despite this region being naturally the most densely populated with intraepithelial macrophages. Nonetheless, a significant reduction in epithelial height indicated impaired secretory function (**Fig 2b**). In more distal regions, up to 10% of epithelial areas exhibited structural damage, characterized by loss of epithelial integrity, pyknotic epithelial cells, exfoliation of epithelial cells, accumulation of immature germ cells in the epididymal lumen and extravasation of sperm into the interstitial space (**Fig. 2a**, **Fig. 2c**). The loss of epithelial integrity was further confirmed by immunofluorescence stainings for Aquaporin 9 (AQP9) and EPCAM, both of which showed marked epithelial discontinuities (**Fig. 2d and Suppl. Fig. S2b, respectively**). Intriguingly, while the gross morphology appeared to recover towards 30 days after depletion (**Fig. 2a**), both the apical AQP9 layer as well as EPCAM integrity exhibited persistent impairments of the epithelial barrier not observed with Tam or Dtx treatment alone (**Fig. 2e and Suppl. Fig. S2b,c**). Notably, there was no increase in apoptotic cell numbers (cleaved caspase 3 staining, **Suppl. Fig. S2b)** and no detectable change in clear cells numbers (**Fig. 2d**). Tam or Dtx treatment alone did not affect epithelial integrity (**Suppl. Fig. S2c**).

**Figure 2:**
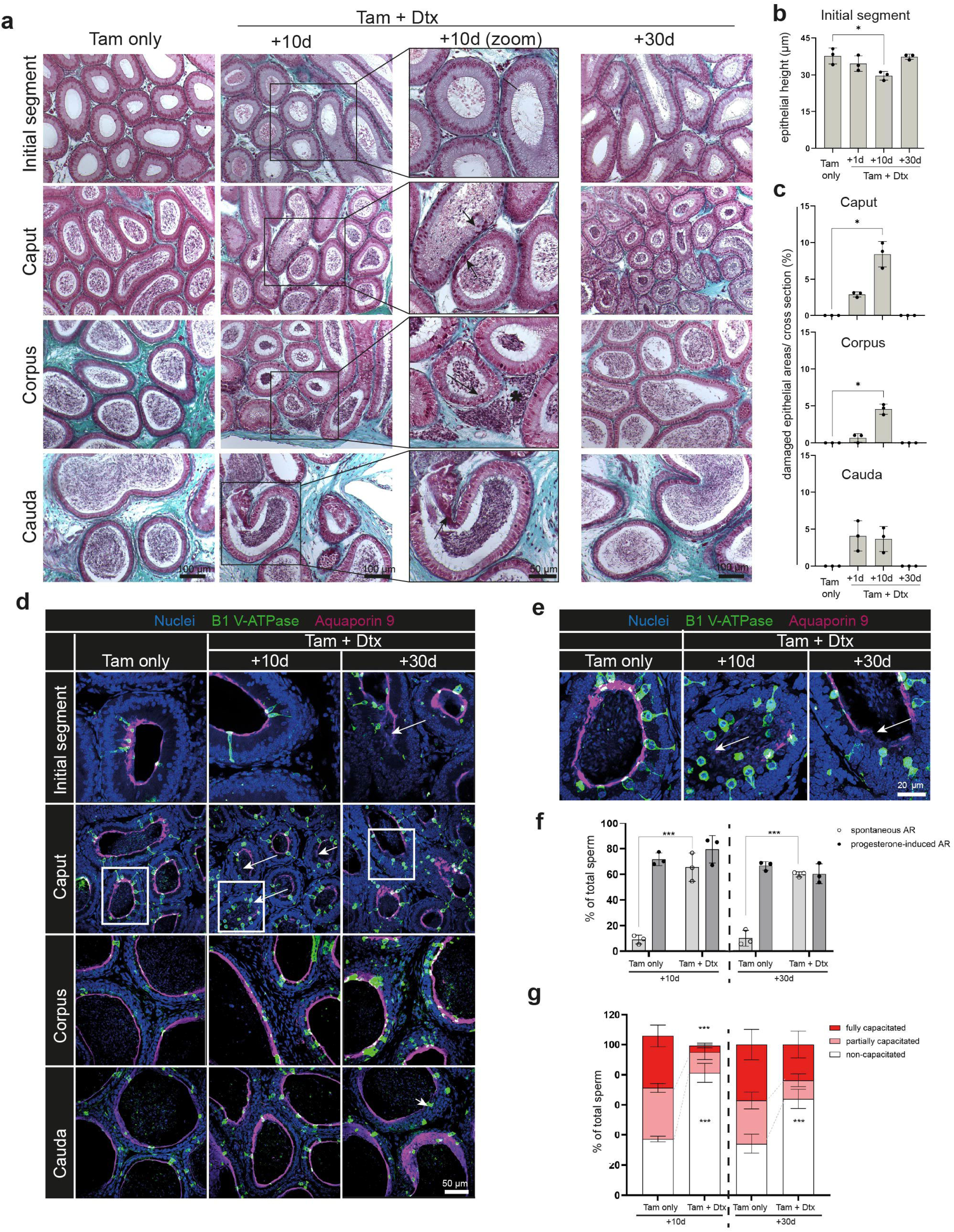
Focal epithelial damage and impaired sperm maturation following Dtx-mediated depletion of CX3CR1^+^ macrophages under physiological conditions. **(a)** Histological analysis of proximal (Initial segment/ Caput) and distal (Corpus/ Cauda) regions at 10 and 30 days post-depletion. Masson Goldner trichrome staining. Arrows denote epithelial disruption, and asterisks indicate extravasated sperm. **(b)** Quantification of epithelial height in the initial segment at the indicated time points. Each data point represents one biological replicate (n=3); 9-10 duct cross-sections (each containing 7-8 epithelial areas) were measured and averaged per sample. **(c)** Frequency of damaged epididymal duct cross-sections in caput, corpus and cauda at indicated time points post-depletion (n=3). **(d, e)** Confocal microscopy of V-ATPase^+^ clear cells (green) and apical aquaporin 9 (purple) layers in IS, caput, corpus, cauda at 10 and 30 days post-depletion. Nuclei are stained with DAPI (blue). Arrows indicate disrupted aquaporin layers. Lined boxes highlight regions enlarged in (E). **(f)** Acrosome reaction (AR) in cauda-derived spermatozoa after in vitro capacitation with and without progesterone stimulation, assessed at 10 and 30 days post-depletion in Tam only and Tam+Dtx-treated mice (n=3 per group). AR was evaluated using PNA staining (see representative images in Fig. S2D). **(g)** Assessment of sperm capacitation by tail protein tyrosine phosphorylation using 4G10 antibody. Sperm were classified as non-capacitated (4G10⁻), partially capacitated (spotted 4G10⁺), or fully capacitated (uniform 4G10⁺), and quantified by confocal microscopy (≥200 sperm per replicate, *n* = 3; representative patterns shown in Fig. S2d). Data are presented as mean ± SD. Statistical analysis was performed using Kruskal– Wallis test with Dunn’s post hoc correction (\**p* < 0.05, \*\**p* < 0.005, \*\*\**p* < 0.001).

To assess functional consequences for sperm maturation, we analyzed sperm isolated from the cauda region. Sperm from CX3CR1^+^ macrophage-depleted mice exhibited a significantly elevated rates of spontaneous acrosome reactions (AR), reaching levels comparable to those induced by progesterone stimulation (**Fig. 2f, Suppl. Fig. S2d**). Regarding capacitation, a prerequisite for acquiring hypermotility within the female reproductive tract, significantly fewer sperm displayed tyrosine phosphorylation (a molecular hallmark of successful capacitation) following *in vitro* stimulation (**Fig. 2g, Suppl. Fig. S2d**). Mice depleted of CX3CR1 macrophages did not develop anti-sperm antibodies after 30 days under physiological conditions (**Suppl. Fig. S2e**), consistent with the absence of inflammatory responses during the 30 day course after Dtx-mediated depletion, suggesting that the depletion of CX3CR1 macrophages alone is not sufficient for the development of anti-sperm autoimmunity.

Collectively, these findings indicate that the loss of epithelial integrity following CX3CR1⁺ macrophage depletion directly compromises sperm maturation, likely due to disrupted epithelial function and subsequent alterations in the luminal microenvironment required for proper sperm development, as opposed to anti-sperm responses.

### Influence of the CX3CR1^+^ macrophage depletion on the testicular macrophage pool

Given that testicular macrophages also express CX3CR1^31^ and CX3CR1^+^ macrophage-depleted mice exhibit increased numbers of immature germ cells in the epididymal lumen, we next examined how the depletion strategy affected the testicular macrophage pool. Overall, no specific macrophage subset appeared to be preferentially tdTomato-labeled or selectively depleted, even when considering the spatial distribution and density of macrophages in relation to the spermatogenic stages, suggesting the transgenic construct targeted both interstitial and peritubular macrophages (**Fig. 3a**). Among all immune cells (CD45^+^), overall numbers were reduced by approximately 20%, with around 40% of immune cells showing tdTomato labeling (**Fig. 3b**). Although Dtx-mediated depletion caused milder reductions in the testis compared to the epididymis **(Fig. 1b**), the number of CX3CR1^+^ macrophages per mg tissue was significantly decreased 1 day after Dtx-treatment, followed by recovery at 10 and 30 days post-depletion, similar to the pattern observed in the epididymis (**Suppl. Fig. S1b, Suppl. Fig. S3a**). When distinguishing between MHC-II^+^ and MHC-II^-^ macrophages, the MHC-II^+^ subset was predominantly tdTomato-labeled and thus selectively depleted by Dtx, whereas MHC-II^-^ macrophages remained largely unaffected (**Fig. 3c, Suppl. Fig. S3b**). High-dimensional analysis, confirmed that tdTomato expression was restricted to CX3CR1^+^ macrophages, which were specifically targeted by the depletion strategy (**Fig. 3d**). Other immune cell populations remained stable and were unaffected by either combined treatment or by Tam or Dtx alone (**Fig. 3e, Suppl. Fig. S3c**).

**Figure 3:**
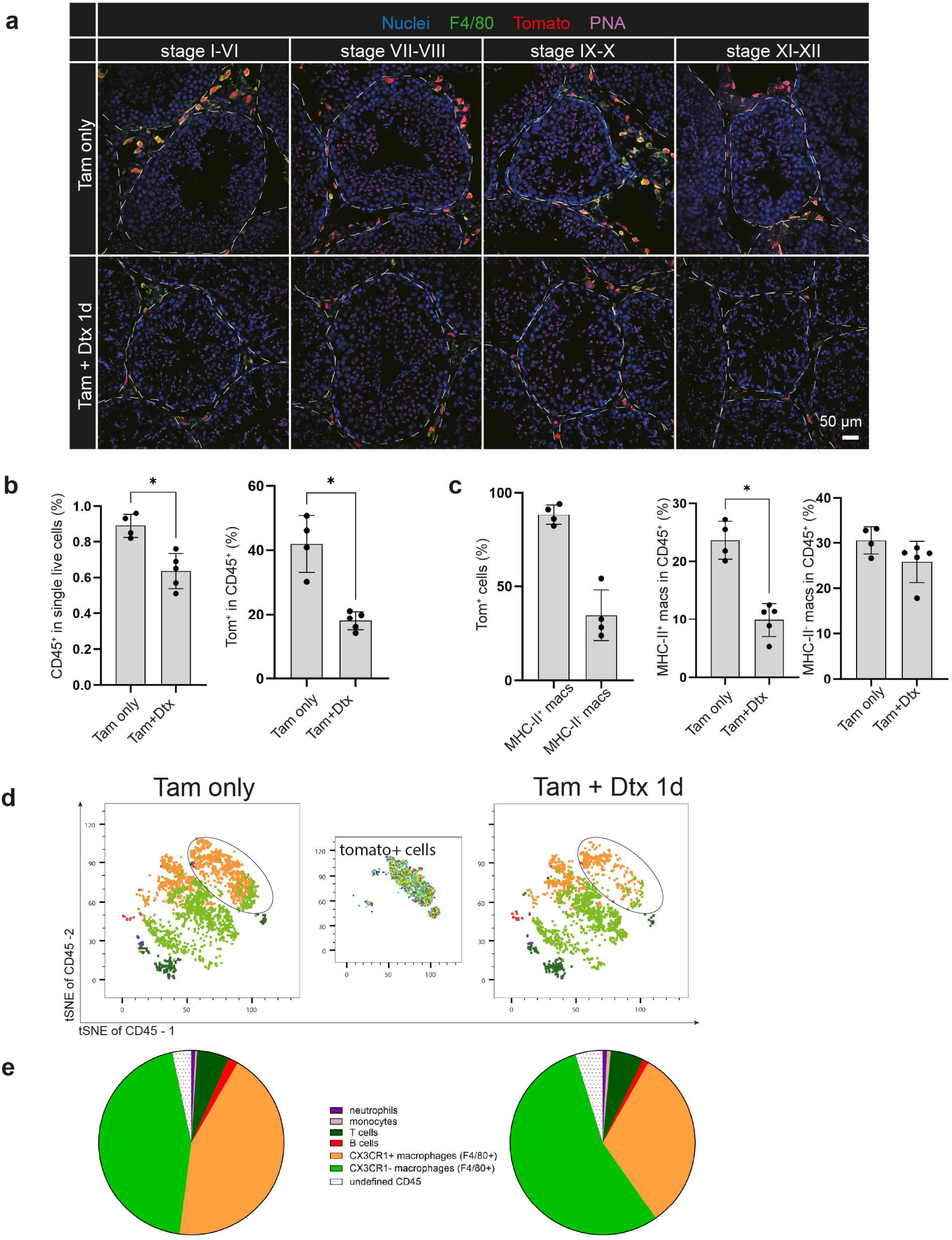
Impact of Dtx-mediated depletion on testicular macrophages at 1 day after depletion. **(a)** Confocal microscopy images showing the distribution of Tom⁺ cells (red) in the testis, aligned with distinct stages of spermatogenesis. Macrophages were co-stained with an antibody against F4/80 (red) and nuclei were stained using DAPI (blue) and developing acrosomes were visualized with PNA (purple) to distinguish spermatogenic stages. **(b)** Relative abundance of CD45⁺ cells among live single cells, and proportion of Tom⁺ cells within the CD45⁺ compartment, one day after depletion in Tam only and Tam+Dtx-treated (+1d) mice (*n* = 4–5 from two independent experiments). **(c)** Proportion of Tom⁺ cells within MHC-II⁺ and MHC-II⁻ testicular macrophage subsets, and relative ratios of MHC-II⁺ and MHC-II⁻ macrophages within total CD45⁺ cells, in Tam only and Tam+Dtx-treated mice (+1d) after depletion (*n* = 4–5). **(d)** t-SNE plots (opt-SNE with iterations: 500, perplexity: 20) of testicular CD45^+^ populations from representative Tam only and Tam+Dtx-treated mice, one day post depletion. CD45^+^ cells were downsampled, gated as shown in Fig. S1E and overlaid onto the t-SNE plots. Tomato+ cells were separately gated and plotted on the t-SNE coordinates. Circles indicate differences in CX3CR1^+^ macrophages between Tam only and Tam+Dtx-treated mice. **(e)** Pie charts showing the relative composition of immune cell populations within the testicular CD45^+^ compartment in Tam only and Tam+Dtx-treated mice, one day post-depletion. Cell populations were defined using the gating strategy in Fig. D1E and are color-matched to the t-SNE plots in (D). Data are presented as mean ± SD from two independent experiments. Statistical analysis was performed using the Mann–Whitney U test (\**p* < 0.05, \*\**p* < 0.005, \*\*\**p* < 0.001).

### Loss of CX3CR1^+^ macrophages results in impaired spermatogenesis and steroidogenesis

Previous studies employing a related model that targets all macrophages, including those originating from CX3CR1⁺ progenitors, demonstrated disrupted spermatogonial differentiation^17^. Building on these findings, we investigated how selective depletion of CX3CR1⁺ macrophages affects spermatogenesis in our model.

Detailed analysis revealed distinct alterations across spermatogenic stages (**Fig. 4b**). Early haploid germ cells (spermatids) were frequently detached from the seminiferous epithelium at stages I–III and IV–VI, and elongated (advanced) spermatids were markedly reduced in macrophage-depleted mice, indicating reduced spermatogenic output. At stage IX, immediately following sperm release, apoptotic cells and debris accumulated at the apical seminiferous epithelium (**Fig. 4c**), as quantified by cleaved caspase–3 immunostaining (representative image in **Suppl. Fig. S4e**). The reduction in elongated (advanced) spermatids was accompanied by increased apoptotic spermatocytes at stage XII – the stage of second meiotic division - suggesting defective germ cell differentiation and impaired meiotic progression (**Fig. 4d**).

**Figure 4:**
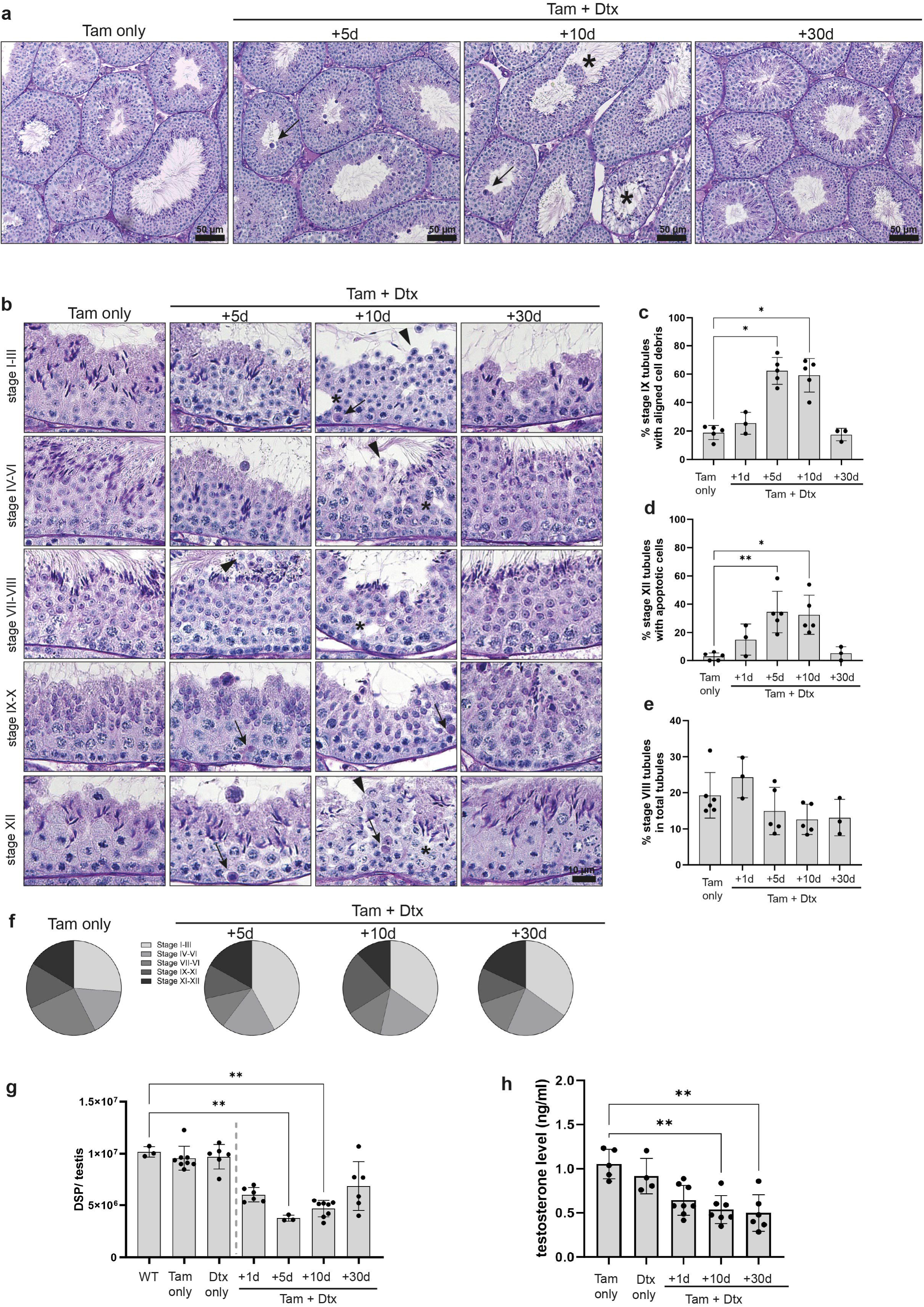
Spermatogenic disruption in the testis following macrophage depletion under physiological conditions. **(a)** Gross histological analysis of testis from Tam only and Tam+Dtx-treated mice at 5, 10 and 30 days post-depletion. Periodic acid-Schiff (PAS)-staining. Arrows indicate round apoptotic cells and asterisks denote focal damage in seminiferous tubules. Periodic acid Schiff’s staining. **(b)** Higher-magnification images highlighting histological abnormalities at specific spermatogenic stages in Tam only and Tam+Dtx-treated mice. Arrowheads indicate exfoliating round spermatids. Arrows mark apoptotic cells and asterisks denote structural lesions in the seminiferous epithelium. **(c)** Proportion of seminiferous tubules with apically aligned cell debris at stage IX (post-spermiation). Each data point represents one biological replicate (n=3-5). **(d)** Proportion of seminiferous tubules showing apoptotic germ cells during stage XII (second meiotic division, each dot represents one biological replicate, n=3-5). **(e)** Frequency of stage VIII tubules (spermiation stage) relative to total tubules (n=3-5). **(f)** Distribution of spermatogenic stages across all tubules, shown as pie charts for Tam only and Tam+Dtx-treated groups (mean of n=3-5). **(g)** Daily sperm production per testis in C57BL/6J wild type, Tam only, Dtx only and Tam+Dtx-treated mice at indicated time points. Each dot represents one biological replicate (n=3-8 from ≥ 2 independent experiments). **(h)** Serum testosterone levels from Tam only, Dtx only and Tam+Dtx-treated mice at indicated time points (n=5-8 from two independent experiments). Data are presented as mean ± SD. Statistical analysis was performed using the Kruskal–Wallis test with Dunn’s post hoc correction (\**p* < 0.05, \*\**p* < 0.005, \*\*\**p* < 0.001).

Although the proportion of tubules at the sperm release stage (stage VII) showed only a non-significant downward trend (**Fig. 4e**), all spermatogenic stages were still present in macrophage-depleted mice at all time points examined (**Fig. 4f**), indicating that spermatogenesis was impaired but not entirely abrogated. By 30 days post-depletion, histopathological changes had reverted to Tam-treated control levels (**Fig. 4a-f**), suggesting that the observed effects were reversible under steady-state conditions. No alterations in gross morphology or spermatogenic staging were observed in mice treated with Tam or Dtx alone (**Suppl. Fig. S4a,b**).

Spermatogenic impairment was reflected in a significant reduction in daily sperm production per testis in macrophage-depleted mice compared to non-induced littermates and single treatment controls (**Fig. 4g**). Despite the quantitative impairment in spermatogenesis, overall total testis weight remained largely unaffected until 30 days after depletion (**Suppl. Fig. S4c**) - at least in relation to total body weight (**Suppl. Fig. S4d**) suggesting a mild rather than severe disruption of overall spermatogenic function.

Notably, steroidogenesis was also impaired, as evidenced by significantly reduced serum testosterone levels (**Fig. 4h**). This decrease had functional consequences, reflected by reduced seminal vesicle size and weight – an organ highly sensitive to testosterone fluctuations (**Suppl. Fig. S4f**).

### Replenished macrophages are distinct from the original population

Given the observed recovery of histopathological alterations in both the epididymis and testis, we investigated by flow cytometry and immunostainings if and how the macrophage pool recovers following Dtx-mediated depletion.

Previously depleted niches were repopulated by macrophages that, at the phenotypic level, resembled those in Tam-only control mice in both proximal and distal epididymal regions as well as the testis 30 days after depletion (**Fig. 5a**). The proportion of F4/80^+^CX3CR1^+^ cells also returned to baseline levels by 30 days post depletion (**Fig. 5b**). Although the macrophage pool quantitatively recovered, high-dimensional analysis and dimensionality reduction based on key subset and activation markers (CX3CR1, CCR2, MHC-II, CD163, and other investigated immune cell markers) revealed that newly recruited macrophages were differentially situated in immune cell clustering when comparing Tam-only and Tam+Dtx-treated mice 30 days after Dtx administration (**Fig. 5c**).We further used high-dimensional flow cytometry to assess differential marker expression of CX3CR1^+^ macrophages in Tam+Dtx and Tam only treated mice at 30 days after depletion. In both the epididymis and testis, MHC-II⁺ macrophages were not only disproportionally affected by the depletion, but also exhibited significantly reduced MHC-II expression after repopulation – particularly in proximal epididymal regions – suggesting diminished antigen-presenting capacity (**Fig. 5d**). Additionally, CX3CR1^+^ macrophages in Tam+Dtx treated mice displayed elevated CCR2 expression (**Fig. 5e**), suggesting a greater contribution of monocyte-derived cells to the repopulating macrophage pool. Under normal steady-state conditions, this feature is restricted to the cauda region^7^.

**Figure 5:**
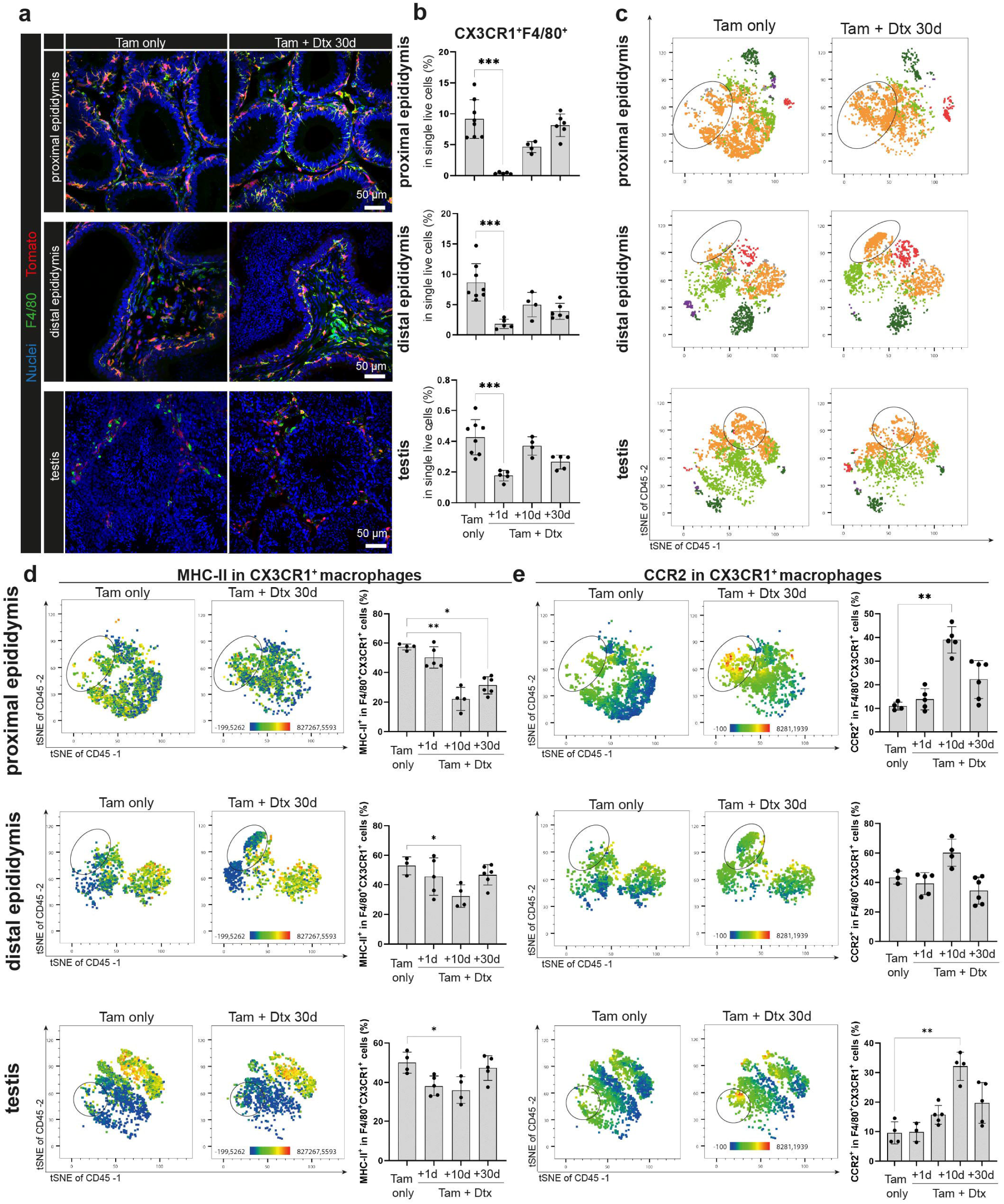
Replenishment of testicular and epididymal macrophages following depletion. **(a)** Confocal microscopy images showing Tom^+^ cells within the proximal and distal regions and the testis in Tam only and Tam+Dtx-treated mice, 30 days post-depletion. **(b)** Proportion of CX3CR1^+^F4/80^+^ macrophages among single live cells at indicated time points post depletion (n=4-8 from two independent experiments). **(c)** t-SNE plots (opt-SNE with iterations: 500, perplexity: 20) of proximal and distal epididymis and testis from representative Tam only and Tam+Dtx-treated mice, 30 days post depletion. CD45^+^ cells were downsampled, gated as shown in Fig. S1E and overlaid onto the t-SNE plots. Circles indicate differences in CX3CR1^+^ macrophages between Tam only and Tam+Dtx-treated mice. **(d, e)** t-SNE plots from C selected for all macrophages with overlaid MHC-II (D) and CCR2 (E) expression and respective proportions of MHC-II^+^ and CCR2^+^ macrophages quantified by flow cytometry at the indicated time points (n=4-8). Circles indicate differences in CX3CR1^+^ macrophages between Tam only and Tam+Dtx-treated mice. Data are presented as mean ± SD. Statistical analysis was performed using the Kruskal–Wallis test with Dunn’s post hoc correction (\**p* < 0.05, \*\**p* < 0.005, \*\*\**p* < 0.001).

The most pronounced differences between repopulated and resident macrophages occurred in proximal epididymal regions, where intraepithelial CX3CR1^+^ macrophages are naturally most abundant and closely associated with the epididymal luminal content.

### Depletion and subsequent dysfunctional repopulation of the CX3CR1^+^ macrophages niche could both contribute to an exacerbated epididymal immune response to bacterial infection

Based on their dense intraepithelial distribution and unique transcriptional profile (characterized by a predominantly homeostatic signature compared to other epididymal macrophage subsets (**Suppl. Fig. S5a**), yet simultaneously adapted (homeostatic in the proximal epididymis and enriched for defense factors in distal regions, **Suppl. Fig. S5b**) to the specific needs of the respective regions) —we hypothesized that intraepithelial CX3CR1⁺ macrophages act as key regulators, tipping the balance in region-specific immune responses toward intraluminal pathogens. To test this, we induced an acute bacterial epididymitis by intravasal injection of uropathogenic *Escherichia coli* (UPEC) following macrophage depletion (**Fig. 6a**).

**Figure 6:**
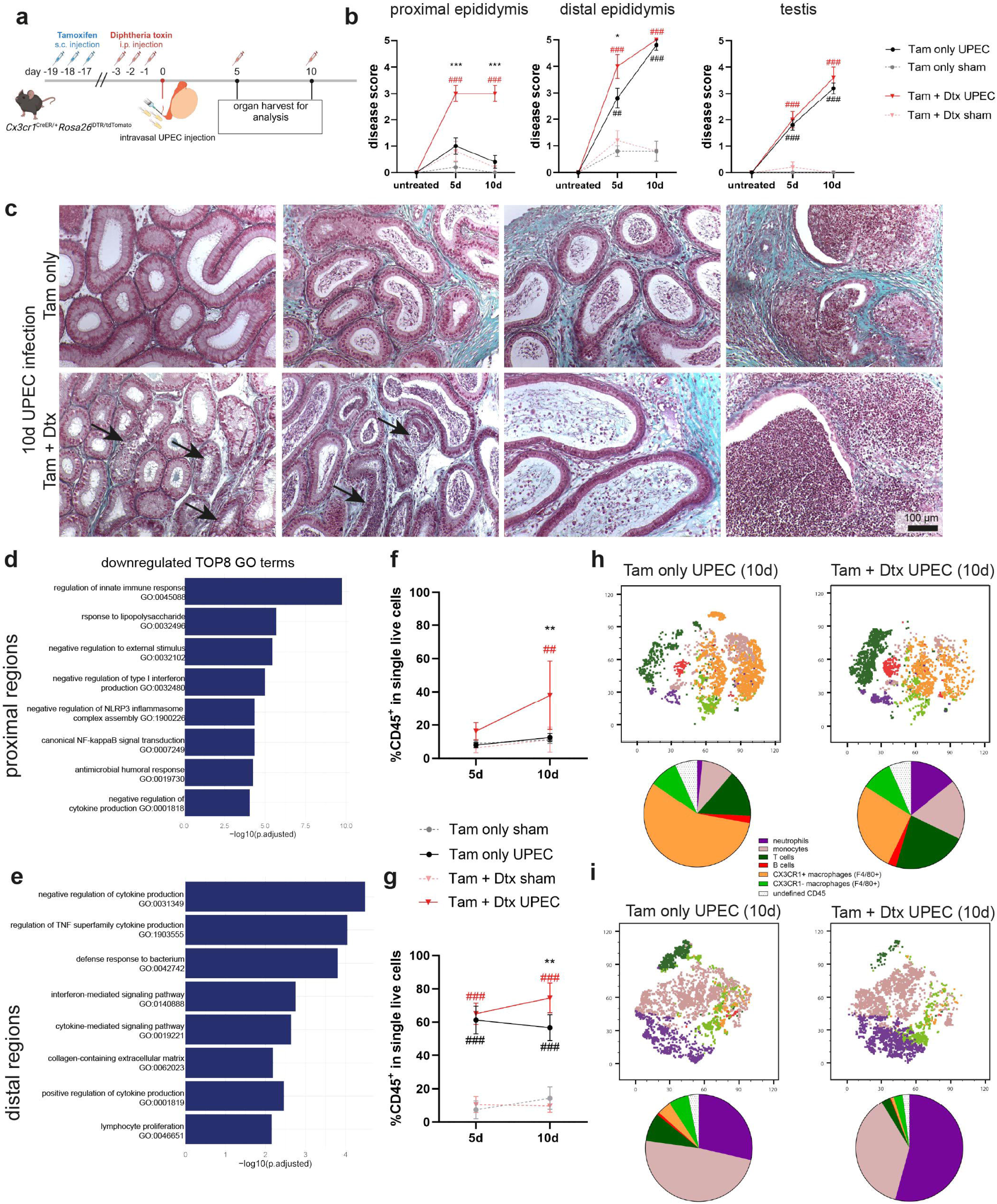
Impact of CX3CR1^+^ macrophage depletion on immune responsiveness during acute bacterial epididymitis. **(a)** Experimental protocol outlining sequential Tam and Dtx treatment in *Cx3cr1*^CreER/+^*Rosa26*^iDTR/tdTomato^ mice, followed by intravasal injection of uropathogenic *Escherichia coli* (UPEC) to induce acute bacterial epididymitis. **(b)** Disease scores in proximal and distal epididymis, and testis at 5 and 10 days after sham or UPEC infection in Tam only and Tam+Dtx-treated mice (n=5 per group from two independent experiments). Scoring was based on established parameters (see Methods). Asterisks indicate the Dtx effect in UPEC-infected mice, red # indicate UPEC effect in Tam+Dtx treated mice and black # indicate UPEC effects in Tam only treated mice. **(c)** Representative histological images from Tam only and Tam+Dtx-treated mice, 10 days post-UPEC infection, showing epithelial damage across all four epididymal regions. Arrows highlight areas of pronounced damage in the IS and caput of Tam+Dtx-treated mice. Masson-Goldner-Trichrome-Staining. **(d, e)** Gene set enrichment analysis (GSEA) based on differentially expressed genes (DEGs) between Tam only and Tam+Dtx-treated mice at 5 days after UPEC infection. Top eight downregulated gene ontology (GO-term shown; (n=3). **(f, g)** Relative abundance of CD45^+^ cells among single live cells at 5 and 10 days after UPEC infection in Tam only and Tam+Dtx-treated mice. Asterisks indicate the Dtx effect in UPEC-infected mice, red # indicate UPEC effect in Tam+Dtx treated mice and black # indicate UPEC effects in Tam only treated mice. **(h, i)** t-SNE plots (opt-SNE with iterations: 1000, perplexity: 30) of CD45^+^ populations within proximal (H) and distal (I) epididymis from representative Tam only and Tam+Dtx-treated mice, 10 days after UPEC infection. CD45^+^ cells were downsampled, gated as shown in Fig. S1E and overlaid onto the t-SNE plots. **(j)** Pie charts showing the relative composition of CD45⁺ immune subsets, defined by the gating strategy in Fig. S1E and color-matched to the t-SNE plots. Data are presented as mean ± SD. Statistical analysis was performed using ANOVA (*p < 0.05, **p < 0.005, ***p < 0.001).

At day 10 after infection, mice depleted of CX3CR1⁺ intraepithelial macrophages exhibited markedly heightened immune response, particularly in proximal epididymal regions that normally remain largely unresponsive when infected with UPEC under control conditions (**Fig. 6b**). The exacerbated immune response was characterized by severe epithelial damage, exfoliation of epithelial cells, interstitial immune infiltrates and extravasation of spermatozoa. Moreover, inflammation also arose earlier across all epididymal regions (**Suppl. Fig. S5c**) with significantly more severe tissue damage, especially in the cauda region, detected by day 5 after infection. In contrast, testicular responses were comparable between groups (**Fig. 6b**).

Transcriptional profiling revealed a pronounced downregulation of genes involved in innate immune regulation, lipopolysaccharide responses, negative regulation of type I interferon production, NLRP3 inflammasome activation and responses to external stimuli in proximal regions in CX3CR1^+^ macrophage-depleted mice. These findings suggest that intraepithelial macrophages normally restrain excessive proinflammatory responses following bacterial infection and thereby preserve tissue integrity (**Fig. 6d**, **Suppl. Fig. S5d**). In distal regions, genes associated with cytokine regulation (e.g. TNFα production), antibacterial defense, and collagen-containing extracellular matrix production were downregulated in CX3CR1^+^ macrophage-depleted mice, indicating a dysregulation of inflammatory processes (**Fig. 6e**, **Suppl. Fig. S5d),** potentially contributing to the observed earlier onset of severe tissue-damaging inflammation.

Consistent with these transcriptional changes seen at day 5 post-infection, significant differences in the magnitude of immune cell infiltration were observed at day 10 in both proximal and distal epididymal regions (**Fig. 6f,g**, respectively). Neutrophils were particularly increased in both regions (**Fig. 6h,i**), while proximal regions additionally showed a significantly amplified ratio of T cells, indicating an activation of adaptive immunity (**Suppl. Fig. S6d**). Interestingly, CX3CR1⁺ macrophages repopulated more rapidly in proximal epididymal regions of UPEC-infected mice and overlap phenotypically with monocytes (**Fig. 6h,i, Suppl. Fig, S6d**), suggesting that infection drives enhanced monocyte recruitment and repopulation of the depleted niches.

Notably, the heightened inflammatory response observed across all epididymal regions was not due to impaired bacterial clearance, as bacterial loads remained comparable between macrophage-depleted and Tam-only control mice at 10 days post-infection (**Suppl. Fig. S6e**).

Within the testis, no significant differences in immune response were observed, as indicated by comparable magnitudes of histopathological alterations between macrophage-depleted and control mice (**Suppl. Fig. S7a,b**). Only minor transcriptional changes occurred, including upregulation of genes involved in extracellular matrix remodeling in CX3CR1^+^ macrophage-depleted mice. Conversely, downregulated processes related to sister chromatid segregation, regulation of chromosome organization, and microtubule dynamics likely reflect the impact of macrophage depletion on germ cell differentiation effects previously noted under uninfected conditions and now potentially exacerbated in the context of inflammation (**Suppl. Fig. S7c**). Consistent with the histopathology, both macrophage-depleted and control groups exhibited only mild immune cell infiltration following UPEC infection, with a trend toward increased neutrophil and T cell accumulation in macrophage-depleted mice (**Suppl. Fig. S7d,e).**

Overall, these findings indicate that CX3CR1⁺ macrophages are critical regulators of immune responses in the epididymis, but to a lesser extent in the testis. In the epididymis they likely serve as key drivers of a tightly controlled immune environment maintaining the delicate balance between effective pathogen defense and preservation of tissue integrity.

## Discussion

Immune regulation and tissue integrity are essential for the specialized microenvironments that support male germ cell development, sperm maturation and fertility. In line with previous studies^11^, our data show that epithelium-associated CX3CR1^+^ macrophages in the epididymis are highly regionally adapted and central to tissue homeostasis in both steady-state and pathological contexts. Although phenotypically distinct from testicular macrophages ^6^, they share the overarching function of supporting key physiological processes. Targeted depletion of CX3CR1^+^-macrophages disrupted testicular germ cell development and epididymal sperm maturation, underscoring their essential role.

Unlike the immune-privileged testis, the epididymis is not a primarily immune-privileged site. Its immune regulation relies heavily on the epithelial integrity (maintained by the blood-epididymis barrier formed by apical junctional complexes between principal cells) and a unique epithelial cell composition^32,33^. Intraepithelial mononuclear phagocytes and epithelial cells, particularly clear cells, are strategically positioned and interact to shape an immune-protective environment in which sperm mature and are stored, shielded from autoimmune responses and pathogens^7,34^. Recent findings suggest that these interactions are region-specific^35^. Although clear cell abundance and morphology remained unchanged, their function is likely impaired following the selective depletion of intraepithelial macrophages, which may contribute to the observed defects in sperm maturation. Indeed, previous studies have suggested that macrophages stimulate clear cell activity (e.g. by the chemokine RANTES) required for luminal acidification under steady-state conditions^16^.

Evidence from efferent duct ligation (EDL) further supports the role of epididymal macrophages in maintaining epithelial homeostasis. EDL triggers apoptosis of epithelial cells by blocking luminal testicular factors. Damaged cells are cleared by efferocytosis^36^, a macrophage-driven process that prevents secondary inflammation and promotes epithelial repair. Moreover, recent studies evidenced that F4/80^+^ macrophages are involved in phagocytosis of extravsated sperm following epithelial damage under steady-state conditions^37^. Together these findings highlight important epithelial cell - macrophage circuits in immunoregulation and maintenance of epididymal tissue homeostasis, functions likely impaired by macrophage depletion and contributing to epithelial disintegration and defective sperm maturation.

The initial trigger for observed long-term changes is unequivocally the depletion of the resident macrophage population, which creates the niche instability that precedes repopulation. However, our data also show that the replenished macrophage pool is phenotypically distinct. Therefore, the persistent impairments in sperm maturation under steady-state as well as the exacerbated immune response to infection - particularly within proximal regions - could be driven not only by the absence of the original regulatory/ homeostatic cells but also by the proinflammatory activity of newly recruited macrophages. This interpretation aligns with recent findings demonstrating that repopulating macrophages exhibit functional and transcriptional profiles distinct from original tissue-resident macrophages, often skewed toward a more proinflammatory phenotype^38,39^.

Of note, macrophage loss caused focal epithelial damage and sperm extravasation without inducing overt inflammation or anti-sperm antibody production under non-infected conditions. Thus, intraepithelial macrophages appear not to be the primary drivers of peripheral tolerance to sperm, a function more typically attributed to dendritic cells^40^ and regulatory T cells^37,41^.

Maintaining epithelial integrity—and thus strict compartmentalization of developing sperm—, however, is crucial to prevent anti-sperm immunity, especially under pathological conditions. In wild type naive mice, the morphology of proximal epididymal regions remains mostly unaffected upon bacterial infection, while distal regions, particularly the cauda, undergo severe tissue remodeling^7–9^. Despite pathogen clearance, chronic inflammation often persists, mirroring clinical epididymitis where up to 40% of patients experience lasting sub-/infertility and develop anti-sperm antibodies^3,4^. Our previous and current studies in experimental mice show that loss of epithelial integrity in the cauda is a critical event, resulting in the leakage of luminal content consisting of millions of sperm with UPEC into the immunologically active interstitial space ^7^ (Pleuger unpublished observations). This exposure may trigger immune cross-reactivity, as pathogens like UPEC can act as adjuvants initiating an immunological cascade that targets other antigens as well^42,43^.

Our findings demonstrate that intraepithelial macrophages influence the onset of inflammation in a region-specific manner and are critical for modulating the course of disease at its initiation, particularly within proximal regions. In doing so, intraepithelial macrophages act in concert with other epithelial cells, including clear cells that were previously demonstrated to actively contribute to the mucosal immunity of the epididymis^34,44^. Among the downregulated genes in proximal epididymal regions of UPEC-infected macrophage-deficient mice, we identified *Cxcl10* – a chemokine with known intrinsic antimicrobial activity against gram-negative bacteria^45^ and immunomodulatory effects via CXCR3 signaling^46^. Notably *Cxcl10* is expressed by clear cells at steady-state in the caput epididymidis, and is further induced upon infection throughout the epididymis^34^. Defining the macrophage-mediated induction/regulation of CXCL10 production in clear cells as well as targeting clear cell–derived CXCL10 and modifying the CXCL10-CXCR3 could provide insights into its role in disease progression and its potential as a prognostic and therapeutic target in translational settings.

In contrast, the macrophage-depleted testis did not show a significantly altered disease course compared to controls. This suggests that partial depletion of CX3CR1^+^ resident macrophages is insufficient to alter the testicular immune response - at least in the context of ascending bacterial infections that primarily affect the epididymis. Notably, even under wild-type conditions, testicular and epididymal immune responses differ, and inflammation-induced tissue damage in the testis is reversible^9^. This likely reflects the immune-privileged status of the testis, where distinct regulatory mechanisms operate compared to the mucosal surface of the epididymis^47^.

Nevertheless, under physiological conditions, we observed clear impairments in both spermatogenesis and steroidogenesis coinciding with the partial loss of testicular macrophages underscoring the sensitivity of testicular function to even partial disruption of macrophage populations and emphasizing the essential role of CX3CR1⁺ macrophages in maintaining tissue homeostasis beyond overt inflammatory responses. Previous studies demonstrated that CX3CR1^+^ progenitors give rise to both interstitial and peritubular macrophages within the testis^31^. In our model, MHC-II^+^ macrophages were predominantly affected, suggesting that mainly, but not exclusively, peritubular macrophages were targeted (though our immunolocalization studies demonstrated a reduction of both peritubular and interstitially located macrophages). Further marker-based stratification of macrophage subsets would be required but was beyond the scope of this study.

Consistent with our observations of spermatogenic impairment, previous studies demonstrated that resident macrophages regulate spermatogonial acitivity indirectly (by interacting with Leydig cells^48,49^ as well as directly through CSF1 and retinoic acid signaling^17,48,49^. For the latter, peritubular macrophage-produced CSF1 targets CSF1R-expressing undifferentiated spermatogonia, facilitated by the close physical association between undifferentiated spermatogonia and peritubular macrophages which are separated only by thin cellular protrusions of the myoid peritubular cells^17,50^.

A related depletion model using *Cx3cr1*^Cre^ (instead of *Cx3cr1*^CreER^), which induced DTX expression in CX3CR1^+^ cells including also other CX3CR1-expressing cells throughout lifetime (including additional immune cell populations) resulted in a more extensive macrophage loss. Minor differences in depletion severity may reflect the tamoxifen-induced DTR expression used in our model, which was designed to selectively target monocyte-independent CX3CR1^+^ macrophages, particularly in the epididymis. Nevertheless, it showed similar phenotypic effects (attributed to disrupted macrophage-spermagonia interactions), and macrophage recovery in the testis^17^. Of note, another study that employed CSF1 antibody-mediated macrophage depletion did not report gross morphological alterations or spermatogenic impairments in the adult testis^51^. Although in our model spermatogenic stages were disrupted, spermatogenesis recovered in parallel with macrophage repopulation, suggesting that testicular function depends on a balanced immune homeostasis.

In both organs, the macrophage niches were replenished over time; however, the phenotype was not fully restored even 30 days after depletion. The increased proportion of CCR2⁺ cells suggests that circulating monocytes contributed to niche repopulation, following an initial phase of proliferation of remaining local macrophage (data not shown). This contrasts with the steady-state in wild-type mice, where testicular and proximal epididymal macrophages are largely maintained by self-renewal, and only distal epididymal regions undergo continuous replenishment by circulating monocytes^6,7^. It is plausible that tissue stress or damage triggers the release of yet unidentified signals that recruit monocytes to repopulate the resident macrophage pool as observed in other tissues^52,53^.

Overall, depletion of CX3CR1⁺ macrophages significantly disrupted immune regulation under both steady-state and inflammatory conditions, particularly in the epididymis, where CX3CR1^hi^ macrophages are integral to mucosal immunity. While systemic effects resulting from testicular dysfunction (particularly regarding steroidogenesis) cannot be excluded, the focal, segment-specific nature of the epithelial damage observed in the epididymis suggests that local mechanisms, likely involving macrophage-epithelial interactions, play a significant role. More importantly, 30 days after Dtx-treatment testosterone levels remain decreased while epithelial integrity is reconstituted, as are macrophage numbers. Intriguingly, although germ cell development and maturation were disrupted in both the testis and epididymis, these effects were largely reversible. This suggests that while these processes are highly sensitive to homeostatic imbalance, tissue-intrinsic mechanisms are sufficient to restore local homeostasis, in part by recruiting circulating immune cells to repopulate the niche. Identifying the factors driving this immune restoration remains a key area for future research. Under inflammatory conditions frequently associated with persistent fertility issues in epididymitis patients, intraepithelial macrophages in the epididymis appear to play a central role in modulating the onset of inflammation. As such, they represent promising targets for therapeutic intervention aimed at improving clinical outcomes and preserving male fertility in immunological-based disorders.

## Materials and Methods

### Experimental mice

All mice used in this study were bred and housed under specific pathogen-free conditions in individually ventilated cages with frequent air changes (15/h). Animals were kept under a 12h light/ dark cycle with ambient temperature (22±2°C) and a relative humiditiy of 55±10%. The following strains were obtained from the Jackson Laboratory: *Cx3cr1*^CreER^ (B6.129P2(C)-*Cx3cr1^tm2.1(cre/ERT2)Jung^*/J, JAX #020940^19^), *Rosa26*^iDTR^ (C57BL/6-*Gt(ROSA)26Sor^tm1(HBEGF)Awai^*/J, JAX #007900^20^), and *Rosa26*^tdTomato^ (B6.Cg- *Gt(ROSA)26Sor^tm9(CAG-tdTomato)Hze^*/J, JAX #007909^21^). Mice were maintained according to the supplier’s instructions at the animal facility of the Justus-Liebig University Giessen, Germany. All animal experiments were approved by the local Animal Ethic Committee (Regierungspräsidium Giessen GI20/25 No. G58/2020 and No. G62/2021) and performed in strict accordance with the Guidelines of the Care and Use of Laboratory Animals of the German law for animal welfare and the European legislation for the protection of animals for scientific purposes (2010/63/EU). For euthanasia before organ collection, mice were deeply anaesthetized by inhalation of 4-5% isoflurane, followed by cervical dislocation or terminal heart puncture.

### Depletion of CX3CR1^+^ cells

For experimental depletion of CX3CR1^+^ cells, *Cx3cr1*^CreER^ mice were crossed with *Rosa26*^iDTR^ mice. Compound homozygous males (*Cx3cr1*^CreER/CreER^*Rosa26*^iDTR/iDTR^) were further crossed with *Rosa26*^tdTomato/tdTomato^ females to enable endogenous labeling of DTR-expressing cells upon tamoxifen treatment. To induce recombination, adult male *Cx3cr1*^CreER/+^*Rosa26*^iDTR/tdTomato^ mice (10-12 weeks old) received 125 µg/g body weight Tamoxifen (‘Tam’ [Sigma T5648] - dissolved in corn oil [Sigma C8267] by sonication for 15 min at 37°C. Tamoxifen was administered subcutaneously once daily for three consecutive days. To avoid a co-depletion of short-lived CX3CR1-expressing cells (i.e. neutrophils, monocytes), Diphtheria Toxin (‘Dtx’, [Sigma D0564] - dissolved in sterile PBS) administration started no earlier than 14 days after the final Tam injection. Dtx was delivered by intraperitoneal injection acutely (500 ng daily on three consecutive days) followed by continuous application of 250 ng every fifth day. Mice were sacrificed at various time points after Dtx-mediated depletion for organ collection and downstream analyses. Day 1 after depletion was defined as the first day following the initial three consecutive Dtx injections (**Fig.1a**).

### Induction of acute bacterial epididymitis

Uropathogenic *Escherichia coli* (UPEC) strain CFT073^22^ was cultured as described previously^23^. To trigger acute epididymitis, mice were anaesthetized through ketamine/xylazine narcosis followed by a small incision at the scrotal area to expose the testis-epididymis-complex. After bilateral ligation of the vasa deferentia, 5 µl UPEC suspension (1-2 x 10^5^ colony forming units [CFU]) were injected into the vasa deferentia adjacent to the cauda epididymidis using a Hamilton syringe. Control ‘sham’ mice underwent the same experimental procedure with an intravasal injection of 5 µl sterile 0.9% NaCl instead of bacteria. Mice were sacrificed at day 5 and 10 after infection by isoflurane inhalation and subsequent cervical dislocation. Each experimental group comprised 5-10 mice per time point, and the entire procedure, including downstream analyses, was repeated in at least two independent experiments.

### Determination of CFU

Testes and epididymides (separated into proximal and distal regions) were collected 5d after UPEC infection and homogenized in 250 µl ice-cold sterile PBS. Serial dilutions (10-fold steps) were prepared and spread onto Luria broth (LB) agar plates (10 mg/ml tryptone, 5 mg/ml yeast extract, 10 mg/ml NaCl, 15 mg/ml agar agar, pH7.0) were incubated upside-down at 37°C for 24h, after which CFU were determined and normalized to tissue weight (per my tissue). Pure UPEC cultures and sterile PBS (processed in parallel with tissue homogenates) were used as positive and negative controls, respectively.

### Histological evaluation

Dissected organs were immediately transferred into Bouińs fixative for 6h at room temperature (RT) before standard paraffin embedding. To improve fixative penetration, the testicular capsule (Tunica albuginea) was gently punctured with a 30G needle. Embedded specimens were cut into 5 µm sections, mounted onto Superfrost Plus slides (Thermo Scientific), incubated overnight at 37°C and stored at RT until further processing. Prior to all histological staining, sections were deparaffinized in xylene and rehydrated through a graded ethanol series. Testicular sections were stained with Periodic Acid Schiff’s reagent and hematoxylin. Epididymal section were stained using the Masson-Goldner trichrome method as described previously^7^. Images were acquired using a Leica Thunder Imager and Leica DM750 microscope. Morphometric analyses were performed using ImageJ V1.53a.

### Scoring of histopathological alterations after UPEC infection

To categorize and compare histopathological alterations after UPEC infection, a previously described scoring systems for epididymitis and orchitis^7,24,25^ were adopted with minor modifications (**Table 1**).

**Table 1:**
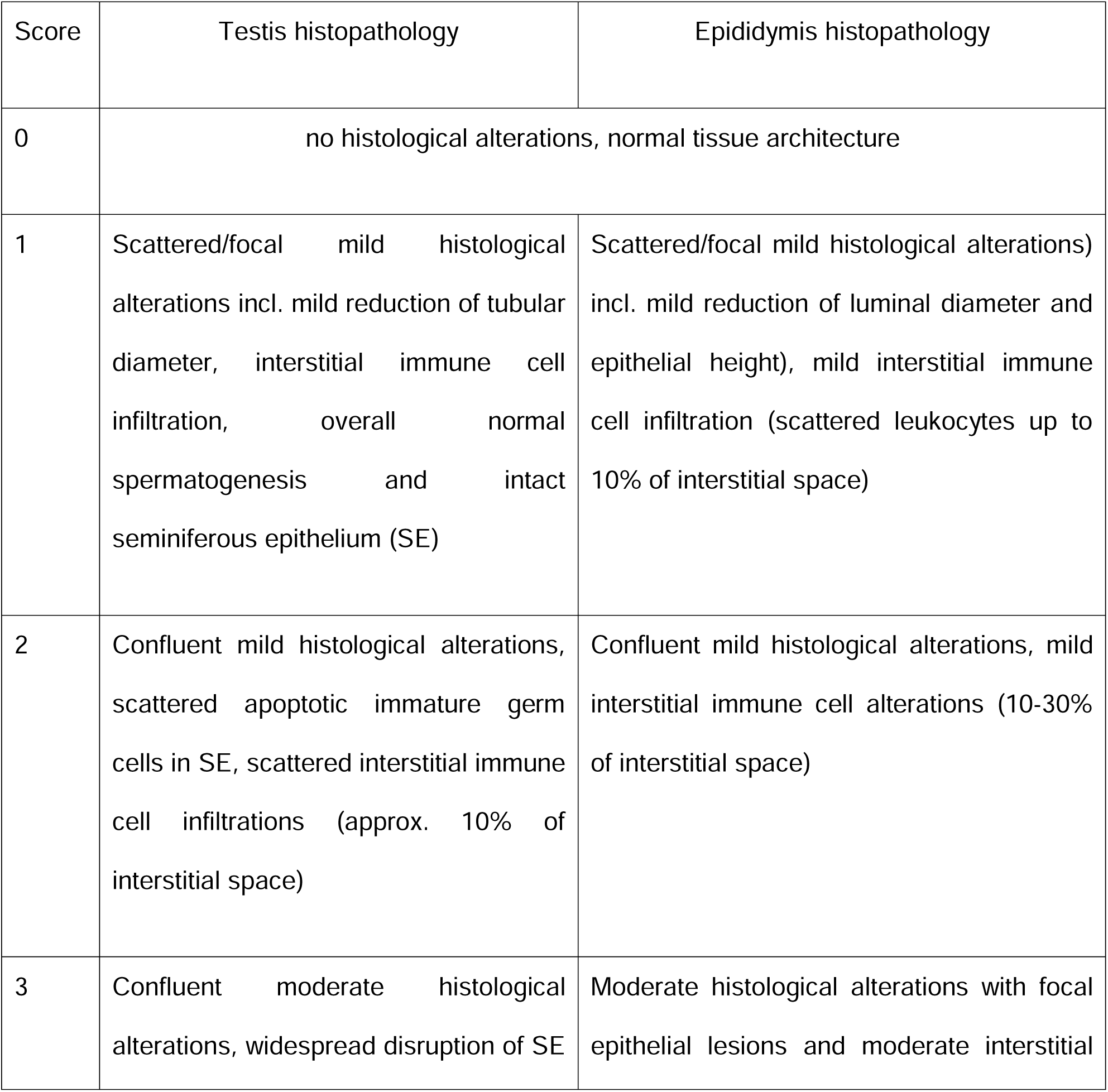
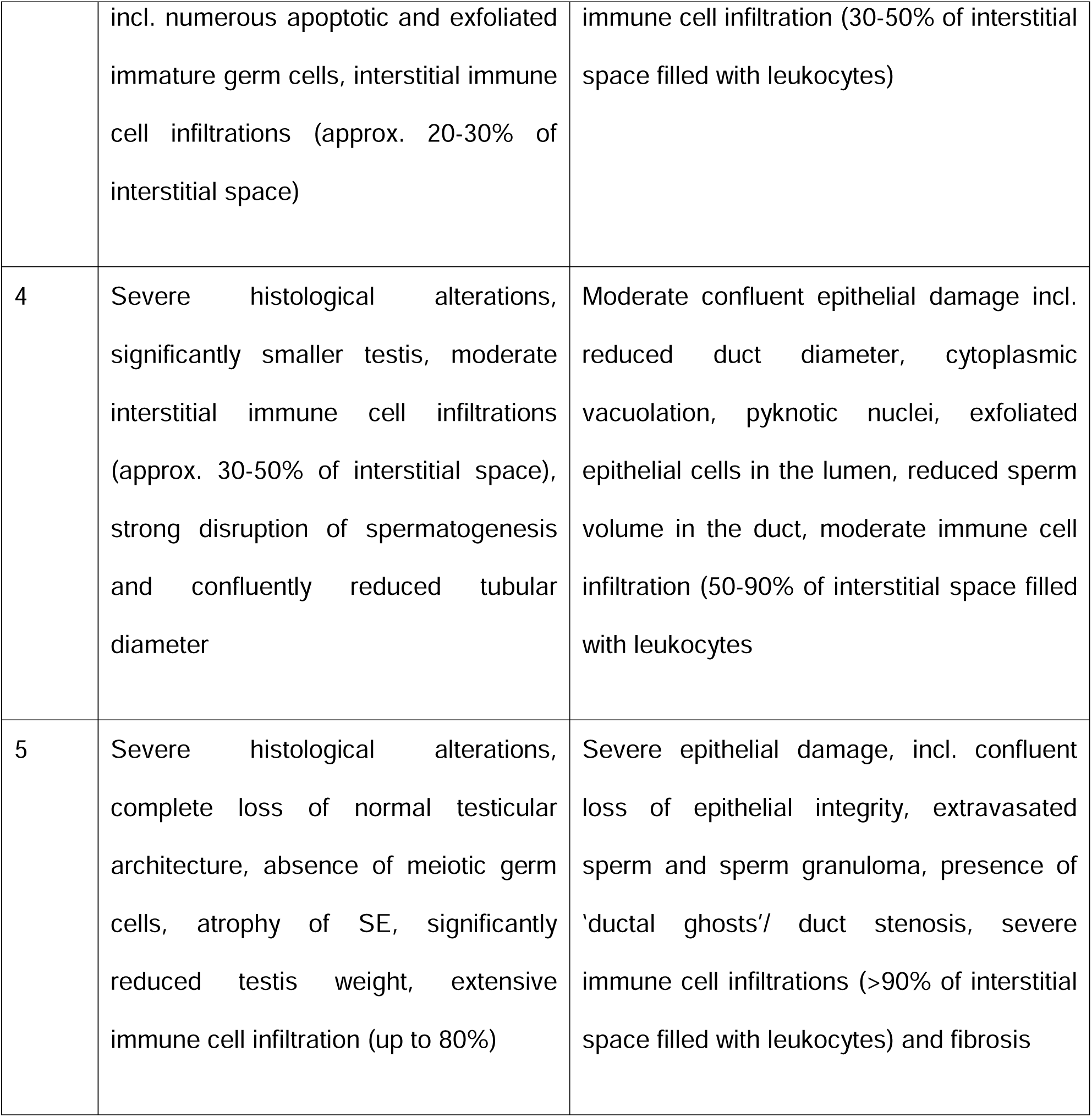
Scoring system of histopathological alterations after UPEC infection.

### Assessment of testicular daily sperm production (DSP)

Decapsulated testes were homogenized in 1 ml DSP buffer (0.9% NaCl, 0.01% Azide, 0.05% Triton-X-100) using a bead-based homogenizer (Retsch). Homogenates were vortexed thoroughly and intact sperm heads counted using a hemocytometer (Buerker chamber 0.0025 mm², depth 0.100 mm). Each homogenate was counted in duplicate, with counts obtained from five of the 16 squares per replicate. DSP was calculated as follows:

(1) Average counted sperm x 50,000 (vol. of counted squares) x (vol. of homogenate [ml] + sample weight [g]) = n_sperm_/ homogenate
(2) n_sperm_/ homogenate/ sample weight [g] = n_sperm_/ g testis
(3) n_sperm_/ g testis x testis weight [g] = n_sperm_/ total testis
(4) n_sperm_/ total testis /4.84 (duration of developing spermatids in step 14-16) = DSP/ testis

### Testosterone measurement

Serum was collected by terminal heart puncture and processed according to standard procedures. Testosterone concentrations were assessed using a commercially available mouse testosterone ELISA kit with respective standardized positive controls (Crystal Chem, **Appendix I**, **Table 2**) following the manufactureŕs recommendations.

### Assessment of anti-sperm antibodies (ASA)

Proteins from untreated wildtype mice were obtained from sperm isolated from the cauda epididymidis by backflushing. Sperm were incubated for 20 minutes on ice in RIPA buffer (supplemented with phosphatase and protease inhibitors), followed by ultrasonication and further 30 min incubation on ice. Samples were centrifuged at 10,000 x g for 20 min, and the resulting supernatant containing soluble proteins was collected to coat high-binding ELISA plates. Antigen solution (200 µl, 6 µg/ml) was incubated overnight at 4°C before washing with PBS-T. Plates were blocked with 3% BSA for 1h at 37°C and then incubated for 2h at 37°C with serum (1:50 in PBS containing 1% BSA) from untreated wildtype, Tam only and Tam+Dtx-treated mice. Uncoated wells treated with serum as well as coated wells treated with PBS served as controls for background reactions.

Bound IgG was detected by incubation with HRP-conjugated goat anti-mouse IgG antibody (0.2 µg/ml) for 2h at RT. Following stringent washing with PBS-T, bound antibodies were visualized by adding TMB substrate for 20 minutes until the desired color developed. Reactions were stopped by adding 100 µl 2M sulfuric acid, and optical density was measured at 450 nm in a microplate reader.

### Functional sperm analysis

Sperm were collected from the cauda epididymidis by backflushing using air pressure to expel sperm from the epididymis^26,27^. Immediately after collection, micropipettes containing sperm were placed in pre-warmed MT6 medium, allowing sperm to swim out for at least 10 min at 37°C. A small fraction of the sperm suspension was collected as the T0 baseline control. Remaining sperm were incubated for 90 min in MT6 medium to induce capacitation. To trigger the acrosome reaction, 15 µM progesterone was added to the capacitation medium for the final 15 min of incubation. Sperm were then collected, washed in PBS and 50 µl aliquots of sperm drops were placed on SuperFrost Plus slides and air-dried overnight. Specimens were post-fixed with 10% neutral-buffered formalin for 15 min, followed by staining of phosphorylated tyrosine using an anti-4G10 antibody (1:1000, **Appendix I Table 2**). Sperm tails were visualized using an anti-acetylated tubulin antibody (1:100, **Appendix I Table 2**). Following overnight incubation with primary antibodies (4°C), corresponding secondary antibodies were applied for 1h at RT in a dark chamber. The acrosome (AR) was visualized with FITC-conjugated peanut agglutinin (PNA, 1 µg/ml) for 30 min at RT. Sperm drops were thoroughly washed and nuclei were stained using DAPI (5 µg/ml). Specimens were mounted using Invitrogen ProLong Gold Antifade Mountant and imaged using a Leica Stellaris 5 confocal microscope. For each biological replicate, at least 200 sperm were evaluated and categorized into (i) fully, partially and non-capacitated sperm according to the 4G10 staining pattern, and (ii) AR-reacted and AR-nonreacted according to the PNA-staining pattern (**Suppl. Fig. S2d**).

### Immunofluorescence

Tissues were fixed in 4% PFA for 6h at 4°C followed by washing in phosphate buffer and overnight incubation in 30% sucrose before embedding in OCT. Sections (20 µm) were air-dried for 20 min at RT in the dark before staining. After washing in TBS+0.05% Tween, pH 7.6 (TBS-T), sections were permeabilized using 0.2% Triton-X-100 and blocked using 3% BSA. Antigen retrieval was performed using either PBS containing 1% SDS (V-ATPase B1 subunit) or 1% SDS+0.1% Triton-X-100 (aquaporin 9) for 4 min. Primary antibodies (Appendix, **Table 2)** were diluted in blocking solution and incubated overnight at 4°C (F4/80 [Bio-Rad]: 10 µg/ml, aquaporin 9 [Battistone lab]: 0.5 µg/ml, V-ATPase B1 subunit [Battistone lab]: 0.3 µg/ml, cleaved-caspase-3 [Cell Signaling]: 1:100 dilution). After thorough washing, corresponding secondary antibodies were applied for 1h at RT in a dark chamber according to manufactureŕs recommendations (**Appendix 1**, **Table 2**). Directly fluorochrome-conjugated antibodies were applied for 1h RT (anti-EpCam-AF594 [BioLegend]: 1 µg/ml). Developing spermatids were visualized by acrosome staining using PNA (FITC-conjugated, 1 µg/ml) applied for 30 min at RT), and nuclei were counterstained using DAPI (5 µg/ml). Specimens were mounted in Mowiol mounting media and imaged using a Leica Stellaris 5 confocal microscope. For quantification, at least 10 fields per regions were counted and averaged per biological replicate.

### Flow cytometry

Before tissue collection, the majority of intravascular CD45^+^ cells were removed by pericardial PBS infusion, as described previously^7^. Epididymides (separated into proximal [segment 1-5] and distal regions [segment 6-10] and testes were collected and immediately transferred into RPMI medium supplemented with 10% FCS. To ensure sufficient cell yield, left and right epididymal fragments from the same individual were pooled to generate a single biological replicate. Tissues were mechanically dissociated by chopping followed by enzymatic digestion for 20 min at 37°C in RPMI containing 10% FCS, 1.5 mg/ml collagenase A and 60 U/ml DNase I. Digested fragments were aspirated through 22G needles four to six times and filtered through a 70 µm cell strainer. After centrifugation at 400xg for 10 min at 4°C, cells were resuspended in RPMI medium with 10% FCS. Non-specific Fc receptor binding was blocked using TrueStain FcX reagent (BioLegend) for 10 min on ice. Cells were stained with antibodies listed in **Appendix Table 2**, following the gating strategy in **Suppl. Fig. S1e** for 30 min at RT in 100 µl of FACS buffer (RPMI medium containing 2 mM EDTA and 0.5% BSA). Controls included omission of the target antibody, incubation with a respective isotype control or fluorescence minus one (FMO) stainings under identical conditions. After antibody incubation, cells were washed twice in FACS buffer. Cell viability was assessed using DRAQ7 staining (1:1000 dilution) for 15 min at RT. Flow cytometry was performed using a BD FACSymphony S6. Data were analyzed using FlowJo v10.10.0 with the DownSampleV3 plugin for expert gating and high-dimensional analysis.

### Statistical analyis

Statistical analysis was performed using GraphPad Prism (version 10.1.2). Detailed information for statistical analysis of particular experiments are stated in the respective figure legend. Overall, a p-value <0.05 was considered significant.

### RNA extraction, library preparation and mRNA sequencing (mRNA-seq)

RNA was extracted from testis, proximal (segment 1-5) and distal epididymis (segment 6-10) using the RNeasy Mini Kit (Qiagen) after bead-based tissue homogenization (Tissue homogenizer Retsch, 2.8 mm stainless steel beads). RNA purification was performed following manufactureŕs recommendation with an additional on-column DNase digestion using RNase-free DNase Set (Qiagen) for 30 min at RT. For genome-wide gene expression analysis, RNA sequencing libraries from isolated mRNA were generated and sequenced by the Institute for Lung Health (ILH), Genomics and Bioinformatics at the Justus-Liebig-University (JLU) Giessen (Germany). For each sample, 1000 ng of total RNA was used for polyadenylated mRNA selection, followed by cDNA sequencing library preparation utilizing the Illumina^®^ Stranded mRNA Prep Kit (Illumina) according to the manufacturer’s instructions. Library quality was assessed by capillary electrophoresis (4200 TapeStation, Agilent) and sequencing of cDNA libraries were performed on the Illumina NovaSeq 6000 platform, generating 50 bp paired-end reads.

Demultiplexing and subsequent FASTQ generation were performed with Illumina’s bcl2fastq (2.19.0.316). Primary read processing (i.e. quality control, filtering, trimming, read alignment and generation of gene specific count tables) was conducted using the nf-core^28^ RNA-seq v3.7 bioinformatics pipeline (NEXTFLOW version 23.04.03) in Docker mode with the *Mus musculus* mm10 reference genome and gene annotation from Illumina’s iGenome repository (https://support.illumina.com/sequencing/sequencing_software/igenome.html). Raw read count tables were imported into R (R Core Team, 2021,^56^) for down-stream processing. Normalization and identification of differentially expressed genes were performed using DESeq2 with default parameter^28^. Gene set enrichment analysis (GSEA) was carried out with the *clusterProfiler* and *fgsea* R packages^29^ using GO and KEGG annotations. Heatmaps were generated using the *complexHeatmap* package^30^. Barplots were generated using the *ggplot2* package. All analysis code is available upon request.

The steady-state transcriptional profiles of *Cx3cr1*^high^ macrophages were compared to (i) other macrophage subsets, and (ii) between proximal (IS, caput) and distal regions (corpus, cauda) by reanalyzing existing data (^7^, GSE208244)).

## Funding and acknowledgement

This work was funded by a project grant to C.P. from the von-Behring-Roentgen-Foundation (grant 69-0029), and honored by the Herbert-Stolzenberg-Award of the JLU to C.P. CP and JUM jointly received funding from the German Research Foundation (DFG) Research Units Programme (FOR) 5644 ‘INFINITE’ project no. 515636567 (CP: PL1017/1-1). The Medical Faculty supported the project by additional intramural funding. JUM was supported by the Rise up! Programme of the Boehringer Ingelheim Foundation (BIS) and the German Research Foundation (DFG) Research Training Group Programme (GRK) 2573/1.

We would like to thank the animal welfare officer and the animal facility team of the JLU Giessen for advisory interactions and expert husbandry of experimental mice. We would also like to thank the Institute of Medical Microbiology of the JLU Giessen for providing a stock of UPEC CFT073.

## Data availability

The transcriptomic datasets generated during this study will be available upon request.

## Author contribution

CP and AM conceived the project and designed experiments. CP, DA and LK conducted experiments (*in vivo* depletion and infection) and collected samples for subsequent phenotypical assessment. CP, DA and LK performed functional sperm analyses, histological analyses, immunostainings and flow cytometry. MSpe and GM provided technical and advisory support for flow cytometry. CP and JUM analyzed flow cytometry data. MAB, MLE and AC provided analytic tools and conducted immunostainings of epididymal epithelial cells. LK and TPK processed samples for sequencing. DA, MB and MSpr performed bioinformatics analyses. CP, AM, SB, MF, JUM and MAB provided scientific advice and conceptual input. CP drafted and wrote the manuscript. All authors were involved in editing and critical revision of the manuscript as well as gave final approval of the submitted and published version.

## Supplemental Figures

**Figure S1: Flow cytometry analysis of immune cell populations in the epididymis under macrophage depletion.**

**(a)** Proportion of CD45^+^ among single live cells, and F4/80^+^CX3CR1^+^ cells among CD45^+^ cells in testis, proximal and distal epididymis of Tam only and Dtx only control mice.

**(b)** Absolute numbers of F4/80^+^CX3CR1^+^ cells per mg tissue in proximal and distal epididymis of Tam only and Tam+Dtx-treated mice at 1, 10 and 30 days post depletion.

**(c)** Distribution of immune cell subsets within the CD45^+^ compartment of Tam only and Dtx only control mice. Immune cells are color-coded as indicated in the legend and defined according to the gating strategy in (E).

**(d)** Proportion of indicated immune cell populations in the CD45^+^ compartment of Tam only and Tam+Dtx-treated mice one day post-depletion. Data are summarized in the pie charts in Fig. 1H-I).

**(e)** Flow cytometry gating strategy to identify indicated immune cell populations under both physiological and pathological conditions in the macrophage depletion model.

Data are presented as mean ± SD from two independent experiments (n=4-5 mice). Statistical analysis was performed using Mann-Whitney *U* test, *p<0.05, **p<0.005, ***p<0.001).

**Figure S2: Histological alterations within the epididymis following macrophage-depletion.**

**(a)** Representative histological images of the IS, caput, corpus and cauda at one day post depletion in Tam only and Tam+Dtx-treated mice (Masson-Goldner-Trichrome-Staining).

**(b)** Confocal immunofluorescence images showing Ep-Cam^+^ epithelial cells (red), macrophages (blue) and cleaved caspase-3 (green) in Tam only and Tam+Dtx-treated mice at indicated time points. Yellow arrows indicate areas with disrupted epithelial integrity. L= Lumen

**(c)** Confocal images of Ep-Cam^+^ epithelial cells (red), macrophages (blue) and cleaved caspase-3 (green) in C57BL/6J, Tam only and Dtx only control mice.

**(d)** Representative images of cauda-derived sperm showing (i) non-, partially- and fully-capacitated states as well as (ii) acrosome-reacted and –nonreacted heads. Tyrosine phosphorylation was detected using an anti-4G10 antibody (red), sperm tails were counterstained with an antibody against acetylated α-tubulin. Acrosomes were labelled with peanut agglutinin (PNA, green), to distinguish acrosome-reacted (-) and non reacted (+) sperm.

**(e)** Detection of IgG against sperm antigens in the serum of wildtype (WT), Tam only and Tam+Dtx-treated mice at 30 days after depletion (n=5 per group from at least 2 independent experiments).

**Figure S3: Flow cytometry analysis of testicular immune cell populations in the macrophage depletion model**

**(a)** Absolute numbers of F4/80^+^CX3CR1^+^ cells per mg tissue in the testis of Tam only and Tam+Dtx-treated mice at 1, 10 and 30 days post depletion.

**(b)** t-SNE plots from Fig 3D selected for all macrophages with overlaid MHC-II expression. Circles indicate differences in CX3CR1^+^ macrophages between Tam only and Tam+Dtx-treated mice.

**(c)** Proportion of indicated immune cell populations in the CD45^+^ compartment of Tam only and Tam+Dtx-treated mice one day post-depletion. Data correspond to pie charts in Fig. 3E).

**(d)** Distribution of immune cell subsets within the CD45^+^ compartment of Tam only and Dtx only control mice. Immune cells are color-coded as indicated in the legend and defined according to the gating strategy shown in Fig. S1E

Data are presented as mean ± SD from two independent experiments (n=4-5 mice).Statistical analysis was performed using Mann-Whitney *U* test, *p<0.05, **p<0.005, ***p<0.001).

**Figure S4: Testicular phenotyping after macrophage depletion.**

**(a)** Representative histological overview images of testes from Tam only and Dtx only control mice, showing no gross abnormalities (Periodic acid-Schiffs staining).

**(b)** Higher-magnification images illustrating normal spermatogenic architecture and no histological alterations at distinct spermatogenic stages in Tam only and Dtx only control mice (Periodic acid-Schiffs staining).

**(c)** Test weight of Tam only, Dtx only and Tam+Dtx-treated mice at indicated time points post depletion (n=3-11).

**(d)** Testis-to-body weight ratio in the same groups and time points as in (C).

**(e)** Representative images of immunohistochemical detection of apoptotic cells using an antibody against cleaved-caspase-3 in testes of Tam only and Tam+Dtx-treated mice at 5 and 10 days post depletion.

**(f)** Seminal vesicle-to-body weight ratio in Tam only, Dtx only and Tam+Dtx-treated mice at 10 and 30 days post depletion. Representative images of seminal vesicles are shown for each group at 30 days.

Data are presented as mean ± SD. Statistical analysis was performed using the Kruskal–Wallis test with Dunn’s post hoc correction (\**p* < 0.05, \*\**p* < 0.005, \*\*\**p* < 0.001).

**Figure S5: Impact of macrophage depletion on epididymal immune responsiveness 5 days post UPEC-infection.**

**(a)** Gene set enrichment analysis based on differentially expressed genes (DEGs) between *Cx3cr1*^high^ macrophages and *Cx3cr1*^low^ macrophages in wildtype steady-state conditions.

**(b)** Gene set enrichment analysis based on differentially expressed genes (DEGs) between *Cx3cr1*^high^ macrophages in proximal and distal epididymal regions in wildtype steady-state conditions.

**(c)** Representative histological images of epididymal regions in UPEC-infected Tam only and Tam+Dtx-treated mice 5 days post infection. Arrows indicate areas with accumulating immune cell infiltrations and epithelial disruption (Masson-Goldner-Trichrome-Staining).

**(d)** Heatmaps displaying downregulated genes in proximal and distal epididymis of Tam only and Tam+Dtx-treated mice. Gene expression data correspond to the downregulated pathways identified in Fig. 6D-E.

**Figure S6: Effects of macrophage depletion on immune responsiveness in sham- and UPEC-infected mice.**

**(a, b)** Representative histological images of epididymal regions ifrom sham-infected Tam only and Tam+Dtx-treated mice at (A) 5 and (B) 10 days after infection Sham mice received intravasal injection of 0.9% NaCl instead of UPEC suspension (Masson-Goldner-Trichrome-Staining).

**(c)** Distribution of immune cell subsets within the CD45^+^ compartment of proximal and distal epididymis as well as testis of sham-infected Tam only and Tam+Dtx-treated mice at 10 days after infection.

**(d)** Relative abundance of indicated immune cell populations in proximal and distal epididymal regions as well as testis of sham and UPEC-infected Tam only and Tam+Dtx-treated mice. Summary data are shown in Fig. 6H,I (epididymis) and S7E (testis).

**(e)** Quantification of bacterial load based on colony-forming units (CFU) per mg tissue in epididymis and testis of Tam only and Tam+Dtx-treated mice 5 days after UPEC infection.

Data are presented as mean ± SD. Statistical analysis was performed using the Kruskal–Wallis test with Dunn’s post hoc correction (\**p* < 0.05, \*\**p* < 0.005, \*\*\**p* < 0.001).

**Figure S7: Impact of macrophage depletion on testicular immune responsiveness following UPEC infection.**

**(a, b)** Testicular histology of sham- and UPEC-infected Tam only and Tam+Dtx-treated mice at (A) 5 and (B) 10 days after infection (Masson-Goldner-Trichrome staining).9

**(c)** Gene set enrichment analysis (GSEA) based on differentially expressed genes (DEGs) between Tam only and Tam+Dtx-treated mice at 5 days after UPEC-infection.

**(d)** Relative abundance of CD45^+^ cells among single live cells at 5 and 10 days after UPEC-infection in Tam only and Tam+Dtx-treated mice.

**(e)** t-SNE plots (opt-SNE with iterations: 1000, perplexity: 30) of testicular CD45^+^ populations from representative Tam only and Tam+Dtx-treated mice, 10 days after UPEC infection. CD45^+^ cells were downsampled, gated as shown in Fig. S1E and overlaid onto the t-SNE plots.

Data are presented as mean ± SD. Statistical analysis was performed using ANOVA (*p < 0.05, **p < 0.005, ***p < 0.001).

## Appendix 1

**Table 2:**
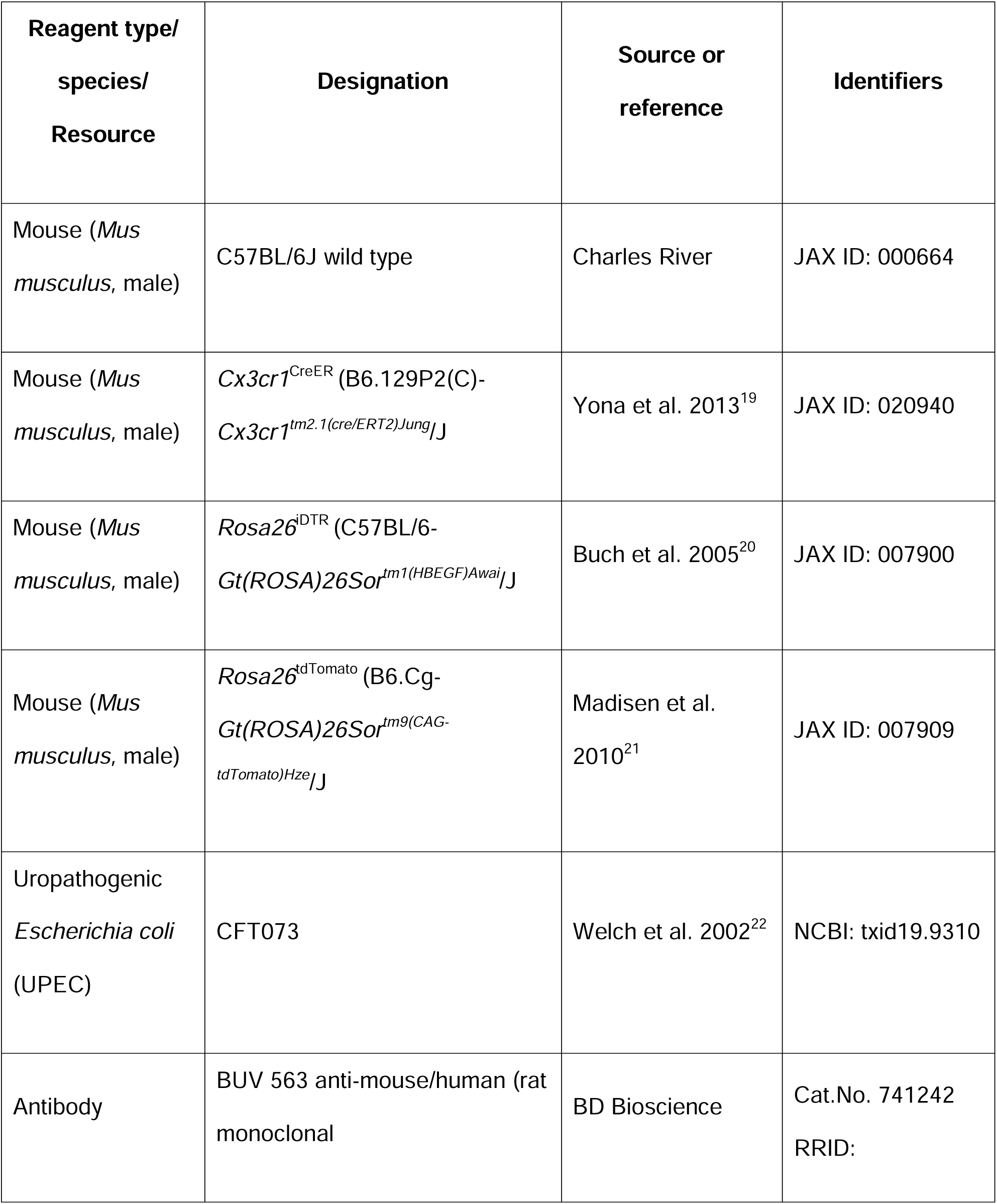
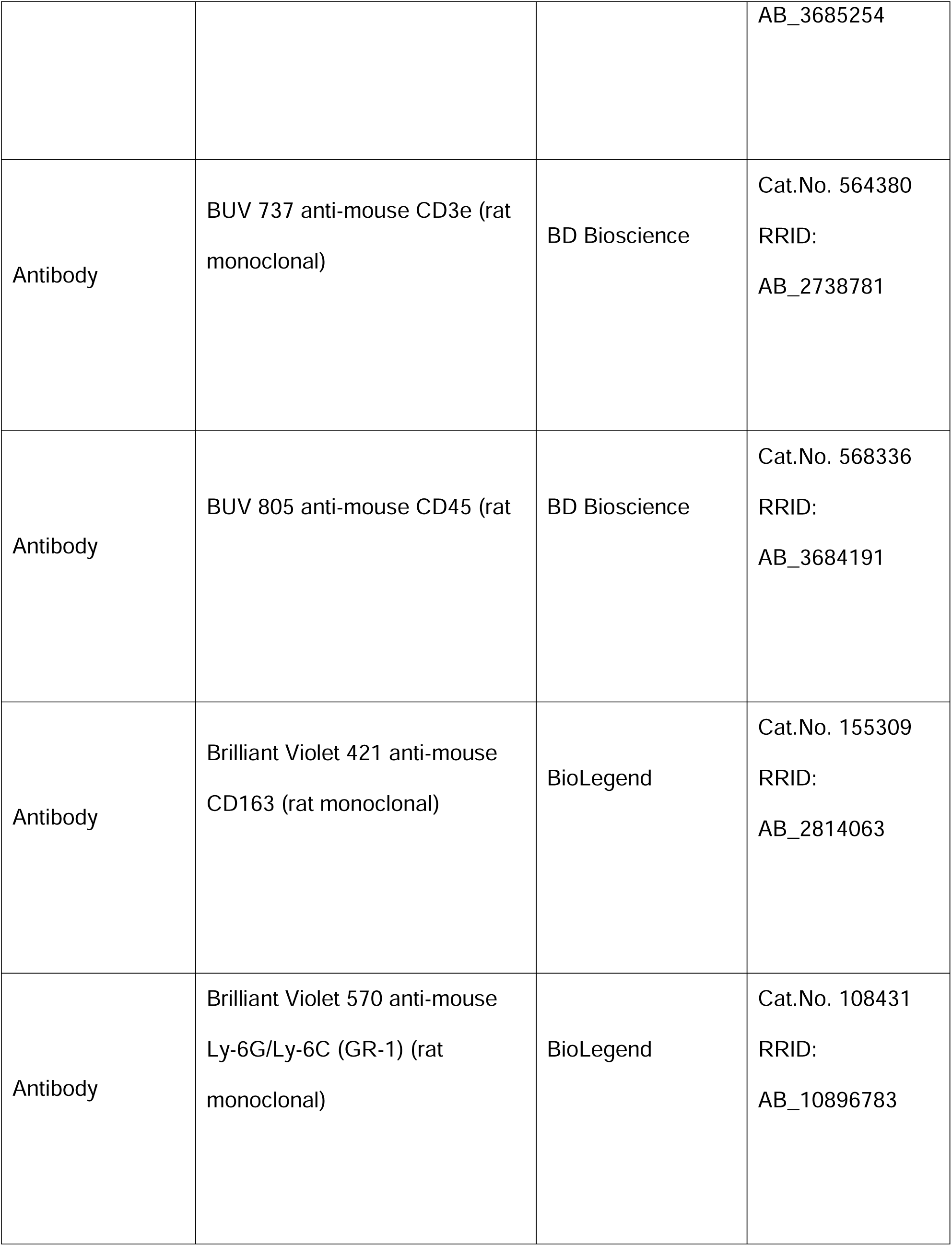
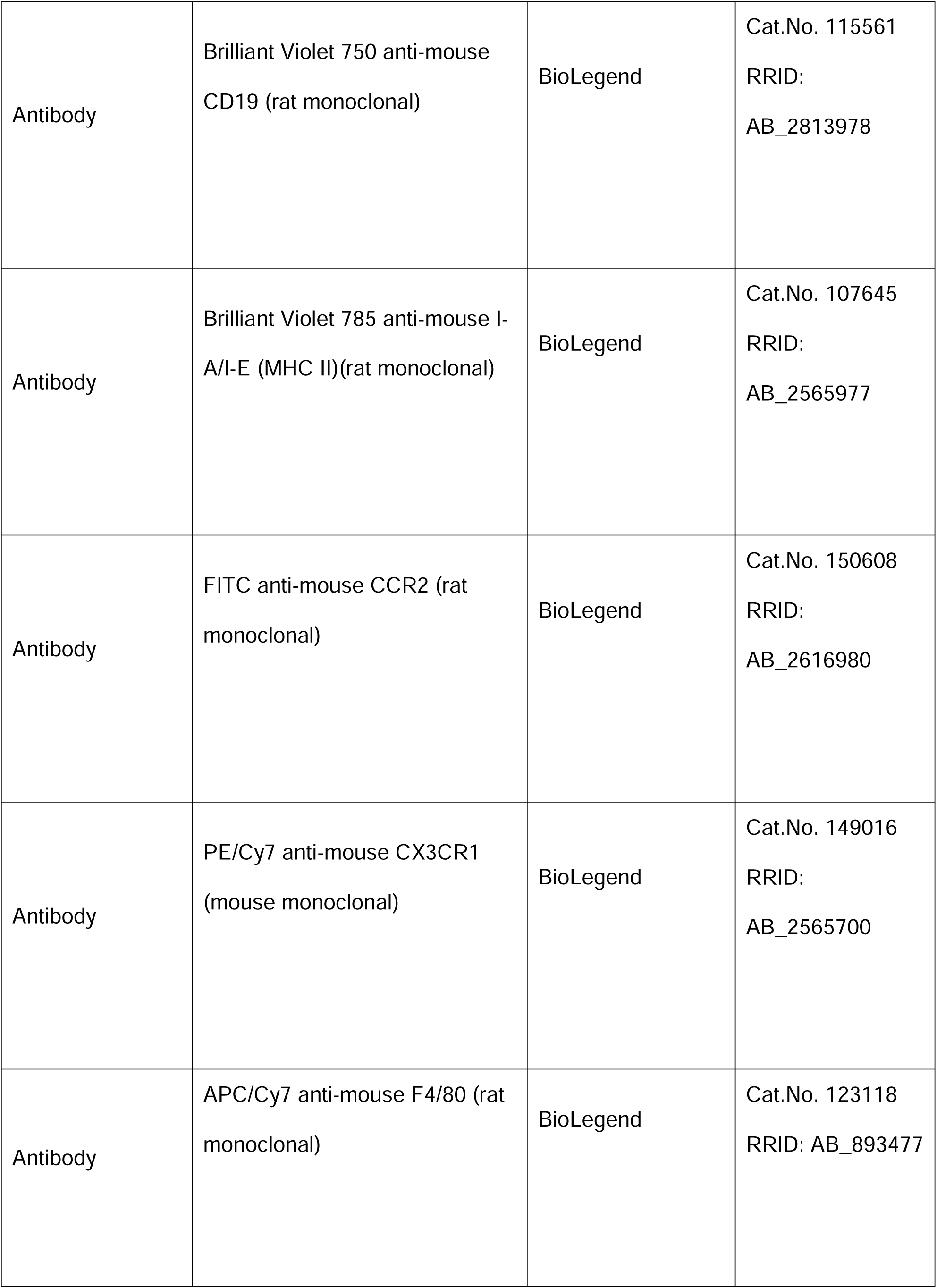
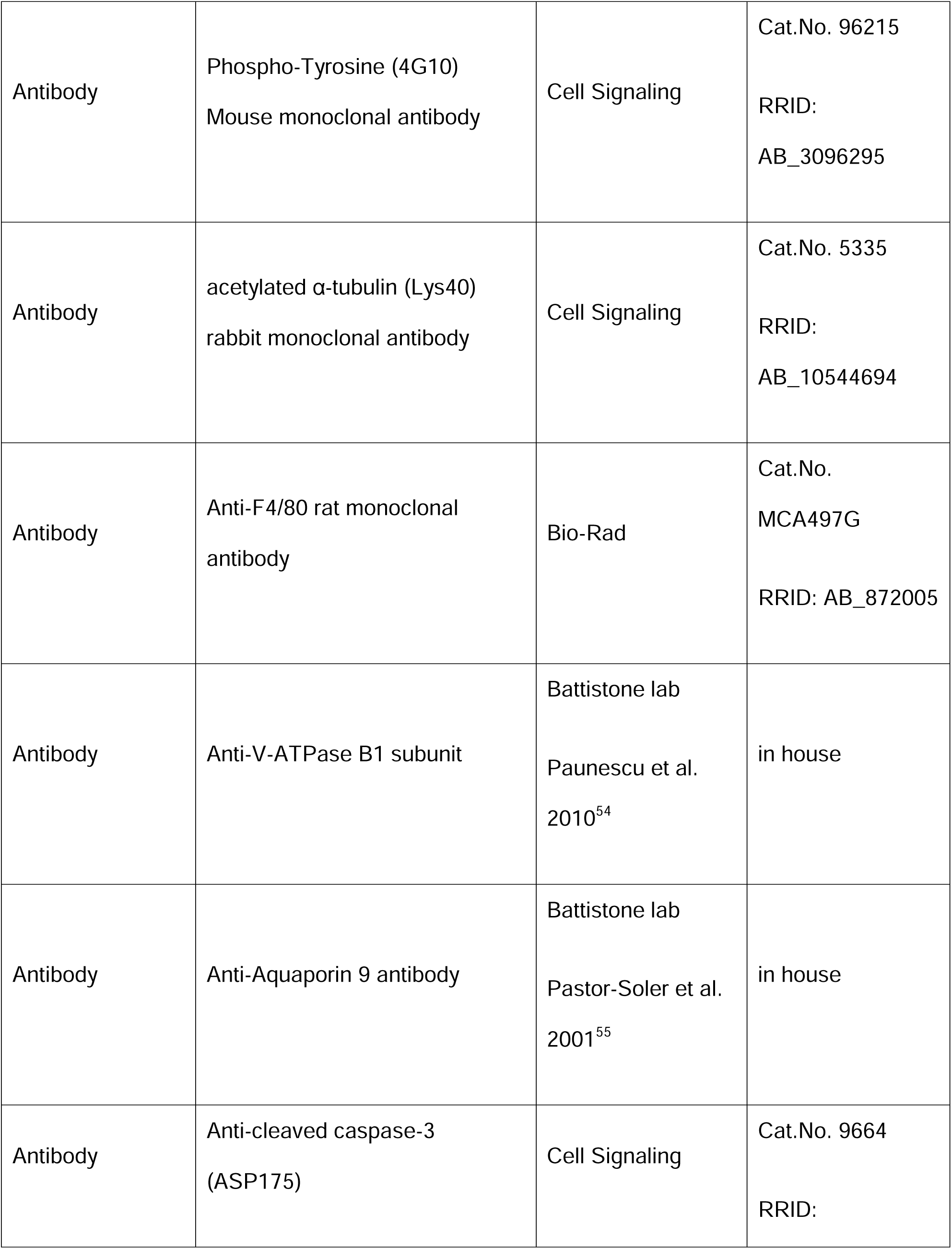
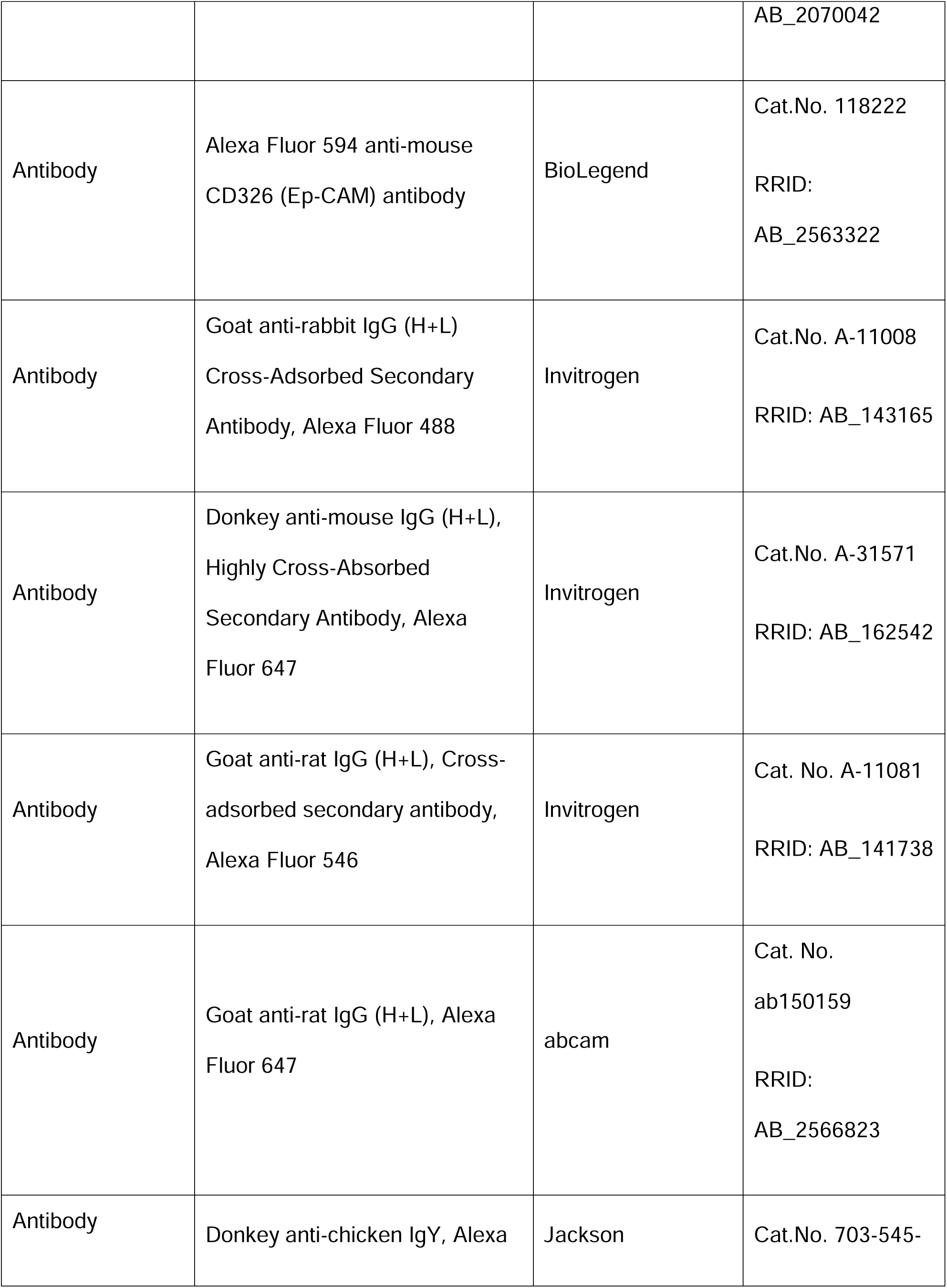
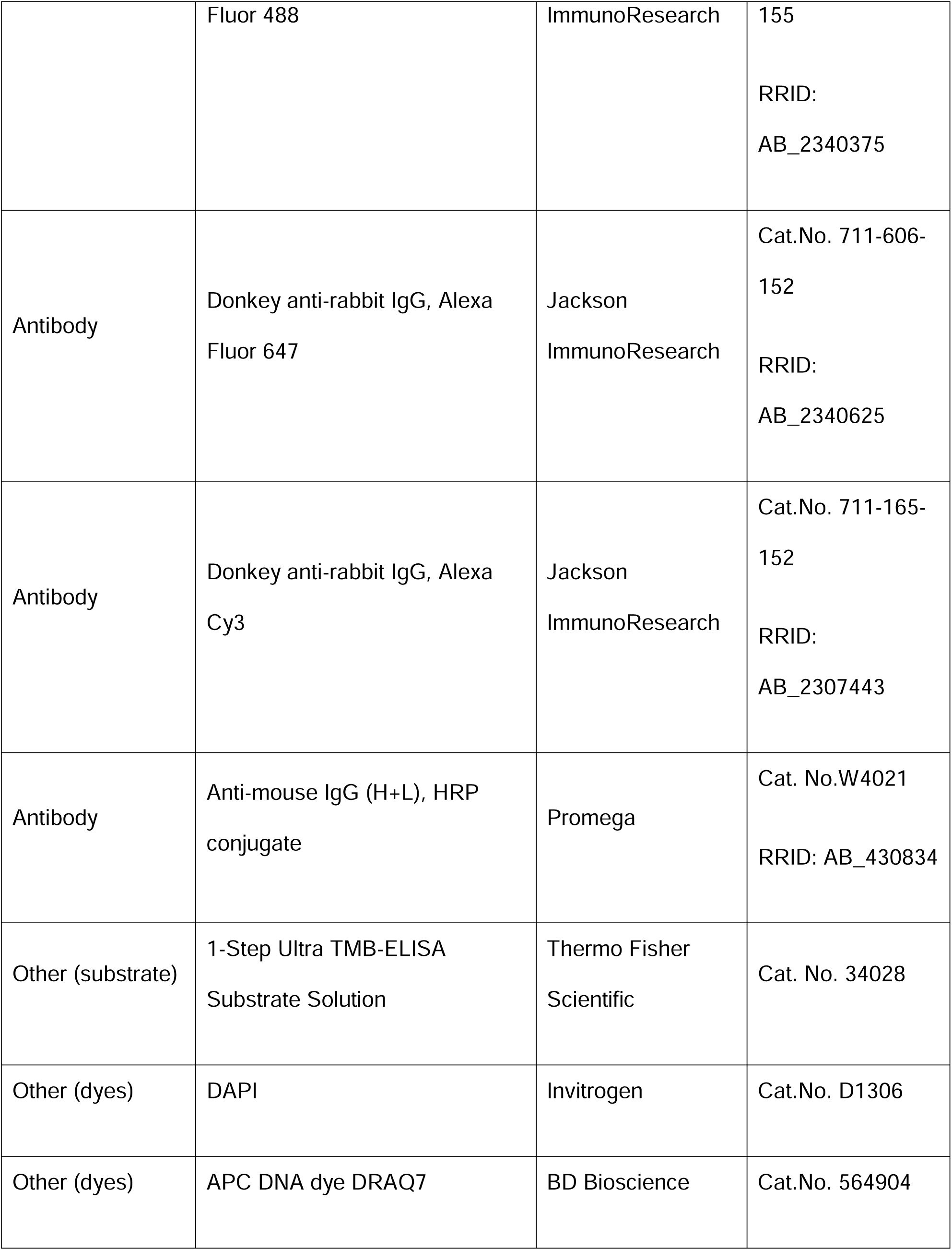
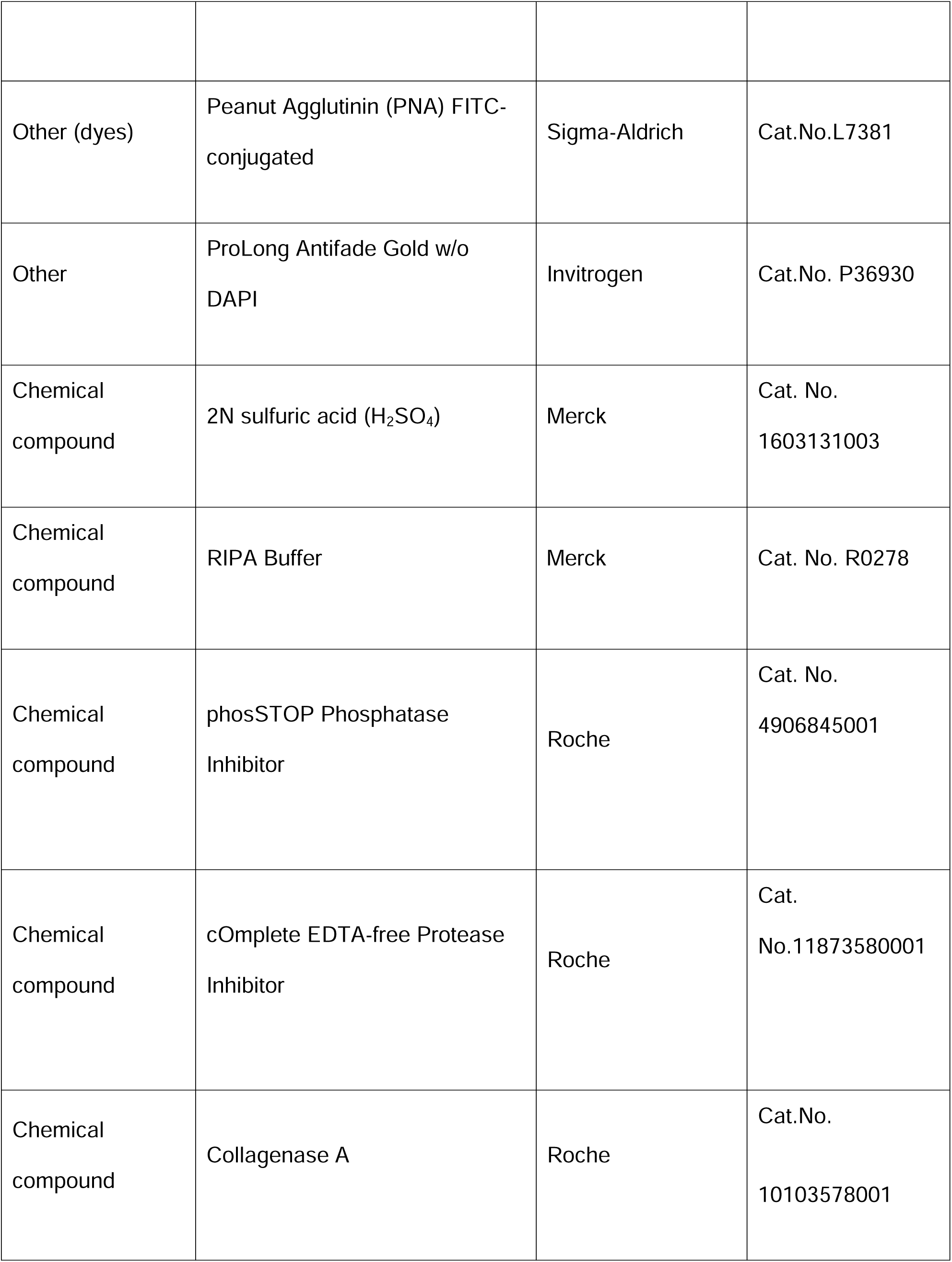
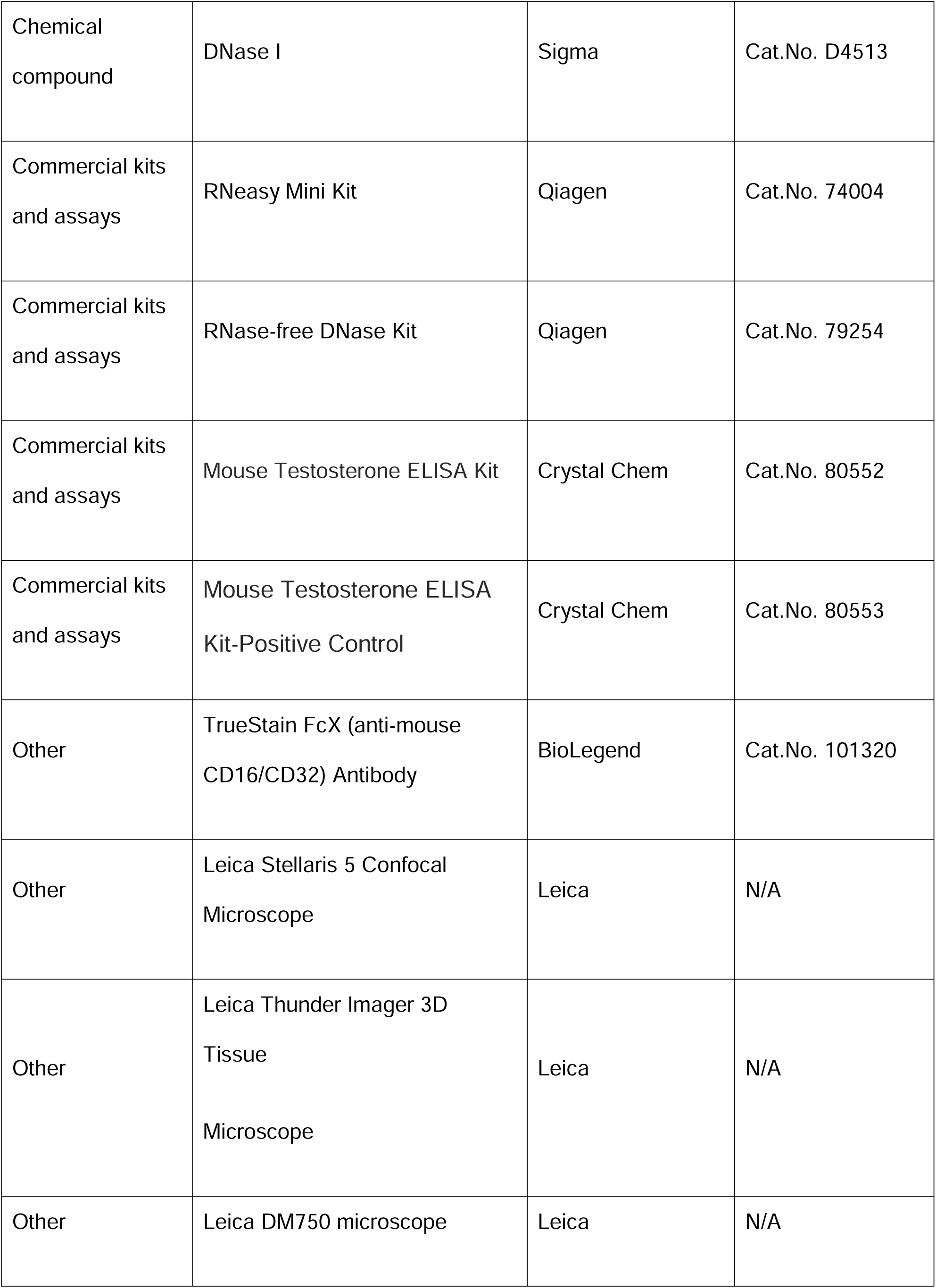
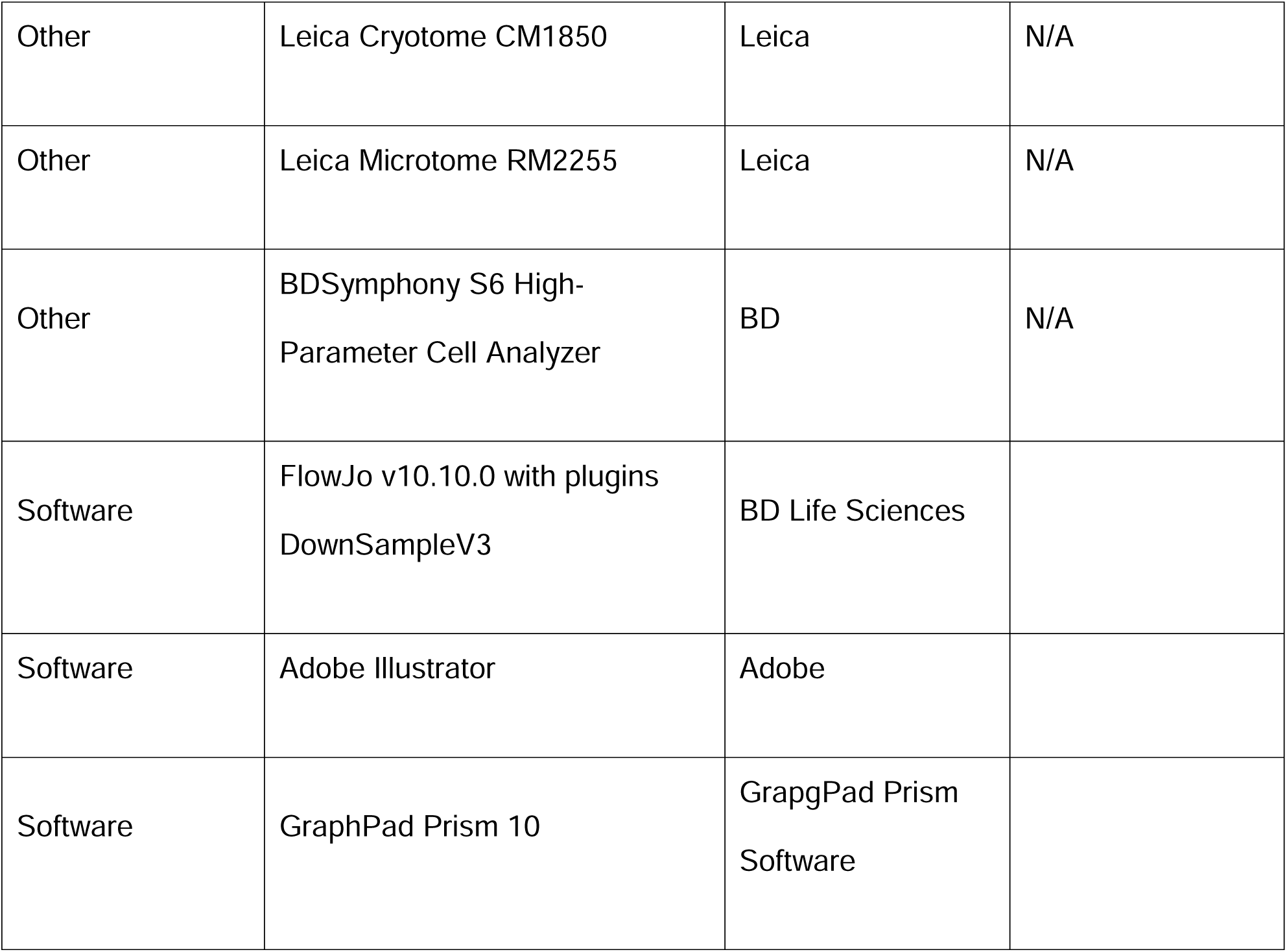
Key resource table.

## References

1. Skerget, S., Rosenow, M. A., Petritis, K. & Karr, T. L. Sperm Proteome Maturation in the Mouse Epididymis. PloS one 10, e0140650; 10.1371/journal.pone.0140650 (2015).

2. Pleuger, C., Silva, E. J. R., Pilatz, A., Bhushan, S. & Meinhardt, A. Differential Immune Response to Infection and Acute Inflammation Along the Epididymis. Frontiers in immunology 11, 599594; 10.3389/fimmu.2020.599594 (2020).

3. Rusz, A. et al. Influence of urogenital infections and inflammation on semen quality and male fertility. World journal of urology 30, 23–30; 10.1007/s00345-011-0726-8 (2012).

4. Lotti, F. et al. Epididymal more than testicular abnormalities are associated with the occurrence of antisperm antibodies as evaluated by the MAR test. Human reproduction (Oxford, England) 33, 1417–1429; 10.1093/humrep/dey235 (2018).

5. Klein, B. et al. Dexamethasone improves therapeutic outcomes in a preclinical bacterial epididymitis mouse model. Human reproduction (Oxford, England) 34, 1195–1205; 10.1093/humrep/dez073 (2019).

6. Wang, M. et al. Two populations of self-maintaining monocyte-independent macrophages exist in adult epididymis and testis. Proceedings of the National Academy of Sciences 118; 10.1073/pnas.2013686117 (2021).

7. Pleuger, C. et al. The regional distribution of resident immune cells shapes distinct immunological environments along the murine epididymis. eLife 11; 10.7554/eLife.82193 (2022).

8. Michel, V. et al. Uropathogenic Escherichia coli causes fibrotic remodelling of the epididymis. The Journal of pathology 240, 15–24; 10.1002/path.4748 (2016).

9. Klein, B. et al. Differential tissue-specific damage caused by bacterial epididymo-orchitis in the mouse. Molecular human reproduction 26, 215–227; 10.1093/molehr/gaaa011 (2020).

10. Voisin, A. et al. Comprehensive overview of murine epididymal mononuclear phagocytes and lymphocytes: Unexpected populations arise. Journal of reproductive immunology 126, 11–17; 10.1016/j.jri.2018.01.003 (2018).

11. Battistone, M. A. et al. Region-specific transcriptomic and functional signatures of mononuclear phagocytes in the epididymis. Molecular human reproduction 26, 14– 29; 10.1093/molehr/gaz059 (2020).

12. Da Silva, N. et al. A dense network of dendritic cells populates the murine epididymis. REPRODUCTION 141, 653–663; 10.1530/REP-10-0493 (2011).

13. Shum, W. W. et al. Epithelial basal cells are distinct from dendritic cells and macrophages in the mouse epididymis. Biology of reproduction 90, 90; 10.1095/biolreprod.113.116681 (2014).

14. Winnall, W. R., Muir, J. A. & Hedger, M. P. Rat resident testicular macrophages have an alternatively activated phenotype and constitutively produce interleukin-10 in vitro. Journal of leukocyte biology 90, 133–143; 10.1189/jlb.1010557 (2011).

15. Bhushan, S. et al. Differential activation of inflammatory pathways in testicular macrophages provides a rationale for their subdued inflammatory capacity. Journal of immunology (Baltimore, Md.: 1950) 194, 5455–5464; 10.4049/jimmunol.1401132 (2015).

16. Feng, X. et al. The Involvement of the Chemokine RANTES in Regulating Luminal Acidification in Rat Epididymis. Frontiers in immunology 11, 583274; 10.3389/fimmu.2020.583274 (2020).

17. DeFalco, T. et al. Macrophages Contribute to the Spermatogonial Niche in the Adult Testis. Cell reports 12, 1107–1119; 10.1016/j.celrep.2015.07.015 (2015).

18. Fijak, M. et al. Infectious, inflammatory and ‘autoimmune’ male factor infertility: how do rodent models inform clinical practice? Human reproduction update 24, 416–441; 10.1093/humupd/dmy009 (2018).

19. Yona, S. et al. Fate mapping reveals origins and dynamics of monocytes and tissue macrophages under homeostasis. Immunity 38, 79–91; 10.1016/j.immuni.2012.12.001 (2013).

20. Buch, T. et al. A Cre-inducible diphtheria toxin receptor mediates cell lineage ablation after toxin administration. Nature methods 2, 419–426; 10.1038/NMETH762 (2005).

21. Madisen, L. et al. A robust and high-throughput Cre reporting and characterization system for the whole mouse brain. Nature neuroscience 13, 133–140; 10.1038/nn.2467 (2010).

22. Welch, R. A. et al. Extensive mosaic structure revealed by the complete genome sequence of uropathogenic Escherichia coli. Proceedings of the National Academy of Sciences of the United States of America 99, 17020–17024; 10.1073/pnas.252529799 (2002).

23. Bhushan, S. et al. Uropathogenic Escherichia coli block MyD88-dependent and activate MyD88-independent signaling pathways in rat testicular cells. Journal of immunology (Baltimore, Md. : 1950) 180, 5537–5547; 10.4049/jimmunol.180.8.5537 (2008).

24. Wijayarathna, R. et al. Region-specific immune responses to autoimmune epididymitis in the murine reproductive tract. Cell and tissue research; 10.1007/s00441-020-03215-8 (2020).

25. Nicolas, N. et al. Induction of experimental autoimmune orchitis in mice: responses to elevated circulating levels of the activin-binding protein, follistatin. REPRODUCTION 154, 293–305; 10.1530/REP-17-0010 (2017).

26. Baker, M. A., Hetherington, L., Weinberg, A. & Velkov, T. Phosphopeptide analysis of rodent epididymal spermatozoa. Journal of visualized experiments : JoVE; 10.3791/51546 (2014).

27. Houston, B. J., Conrad, D. F. & O’Bryan, M. K. A framework for high-resolution phenotyping of candidate male infertility mutants: from human to mouse. Human genetics; 10.1007/s00439-020-02159-x (2020).

28. Ewels, P. A. et al. The nf-core framework for community-curated bioinformatics pipelines. Nature biotechnology 38, 276–278; 10.1038/s41587-020-0439-x (2020).

29. Yu, G., Wang, L.-G., Han, Y. & He, Q.-Y. clusterProfiler: an R package for comparing biological themes among gene clusters. Omics : a journal of integrative biology 16, 284–287; 10.1089/omi.2011.0118 (2012).

30. Gu, Z., Eils, R. & Schlesner, M. Complex heatmaps reveal patterns and correlations in multidimensional genomic data. Bioinformatics (Oxford, England) 32, 2847–2849; 10.1093/bioinformatics/btw313 (2016).

31. Gu, X., Heinrich, A., Li, S.-Y. & DeFalco, T. Testicular macrophages are recruited during a narrow fetal time window and promote organ-specific developmental functions. Nat Commun 14, 1439; 10.1038/s41467-023-37199-0 (2023).

32. Mital, P., Hinton, B. T. & Dufour, J. M. The blood-testis and blood-epididymis barriers are more than just their tight junctions. Biology of reproduction 84, 851–858; 10.1095/biolreprod.110.087452 (2011).

33. Gregory, M. & Cyr, D. G. The blood-epididymis barrier and inflammation. Spermatogenesis 4, e979619; 10.4161/21565562.2014.979619 (2014).

34. Battistone, M. A. et al. Novel role of proton-secreting epithelial cells in sperm maturation and mucosal immunity. Journal of cell science 133; 10.1242/jcs.233239 (2019).

35. Battistone, M. A. et al. Immunoregulatory mechanisms between epithelial clear cells and mononuclear phagocytes in the epididymis. Andrology 12, 949–963; 10.1111/andr.13509 (2024).

36. Smith, T. B. et al. Mononuclear phagocytes rapidly clear apoptotic epithelial cells in the proximal epididymis. Andrology 2, 755–762; 10.1111/j.2047-2927.2014.00251.x (2014).

37. Barrachina, F. et al. Regulatory T cells play a crucial role in maintaining sperm tolerance and male fertility. Proceedings of the National Academy of Sciences 120, e2306797120; 10.1073/pnas.2306797120 (2023).

38. Seifinejad, A. et al. Contrasting contribution of resident and repopulated brain macrophages in sustaining sleep-wake circuitry. Communications biology 8, 1339; 10.1038/s42003-025-08781-7 (2025).

39. Louwe, P. A. et al. Recruited macrophages that colonize the post-inflammatory peritoneal niche convert into functionally divergent resident cells. Nat Commun 12, 1770; 10.1038/s41467-021-21778-0 (2021).

40. Pierucci-Alves, F., Midura-Kiela, M. T., Fleming, S. D., Schultz, B. D. & Kiela, P. R. Transforming Growth Factor Beta Signaling in Dendritic Cells Is Required for Immunotolerance to Sperm in the Epididymis. Frontiers in immunology 9, 1882; 10.3389/fimmu.2018.01882 (2018).

41. Elizagaray, M. L., et al. Chronic inflammation drives epididymal tertiary lymphoid structure formation and autoimmune fertility disorders. bioRxiv : the preprint server for biology; 10.1101/2024.11.12.623224 (2024).

42. Nashan, D. et al. Immuno-competent cells in the murine epididymis following infection with Escherichia coli. International journal of andrology 16, 47–52; 10.1111/j.1365-2605.1993.tb01152.x. (1993).

43. Prior, J. T. et al. Bacterial-Derived Outer Membrane Vesicles are Potent Adjuvants that Drive Humoral and Cellular Immune Responses. Pharmaceutics 13; 10.3390/pharmaceutics13020131 (2021).

44. Da Silva, A. A. S., et al. Proton-Secreting Cells as Drivers of Inflammation and Sperm Dysfunction in LPS-Induced Epididymitis. Function (Oxford, England) 6; 10.1093/function/zqaf023 (2025).

45. Schutte, K. M. et al. Escherichia coli Pyruvate Dehydrogenase Complex Is an Important Component of CXCL10-Mediated Antimicrobial Activity. Infection and immunity 84, 320–328; 10.1128/IAI.00552-15 (2016).

46. Liu, M. et al. CXCL10/IP-10 in infectious diseases pathogenesis and potential therapeutic implications. Cytokine & growth factor reviews 22, 121–130; 10.1016/j.cytogfr.2011.06.001 (2011).

47. Hedger, M. P. Immunophysiology and pathology of inflammation in the testis and epididymis. Journal of andrology 32, 625–640; 10.2164/jandrol.111.012989 (2011).

48. Lukyanenko, Y. O., Chen, J. J. & Hutson, J. C. Production of 25-hydroxycholesterol by testicular macrophages and its effects on Leydig cells. Biology of reproduction 64, 790–796; 10.1095/biolreprod64.3.790 (2001).

49. Hutson, J. C. Physiologic interactions between macrophages and Leydig cells. Experimental biology and medicine (Maywood, N.J.) 231, 1–7; 10.1177/153537020623100101 (2006).

50. Biniwale, S. et al. Regulation of macrophage number and gene transcript levels by activin A and its binding protein, follistatin, in the testes of adult mice. Journal of reproductive immunology 151, 103618; 10.1016/j.jri.2022.103618 (2022).

51. Lokka, E. et al. Generation, localization and functions of macrophages during the development of testis. Nat Commun 11, 4375; 10.1038/s41467-020-18206-0 (2020).

52. Scott, C. L. et al. Bone marrow-derived monocytes give rise to self-renewing and fully differentiated Kupffer cells. Nat Commun 7, 10321; 10.1038/ncomms10321 (2016).

53. van de Laar, L. et al. Yolk Sac Macrophages, Fetal Liver, and Adult Monocytes Can Colonize an Empty Niche and Develop into Functional Tissue-Resident Macrophages. Immunity 44, 755–768; 10.1016/j.immuni.2016.02.017 (2016).

54. Păunescu, T. G. et al. cAMP stimulates apical V-ATPase accumulation, microvillar elongation, and proton extrusion in kidney collecting duct A-intercalated cells. American Journal of Physiology-Renal Physiology 298, F643–54; 10.1152/ajprenal.00584.2009 (2010).

55. Pastor-Soler, N. et al. Aquaporin 9 expression along the male reproductive tract. Biology of reproduction 65, 384–393; 10.1095/biolreprod65.2.384 (2001).

56. R Core Team (2021). R: A language and environment for statistical computing. R Foundation for Statistical Computing, Vienna, Austria. URL https://www.R-project.org/.

