## Supplementary figures and images for "Depletion of CX3CR1^+^ macrophages results in disrupted functionality and immune surveillance within epididymis and testis"

### Supplemental Figure 1

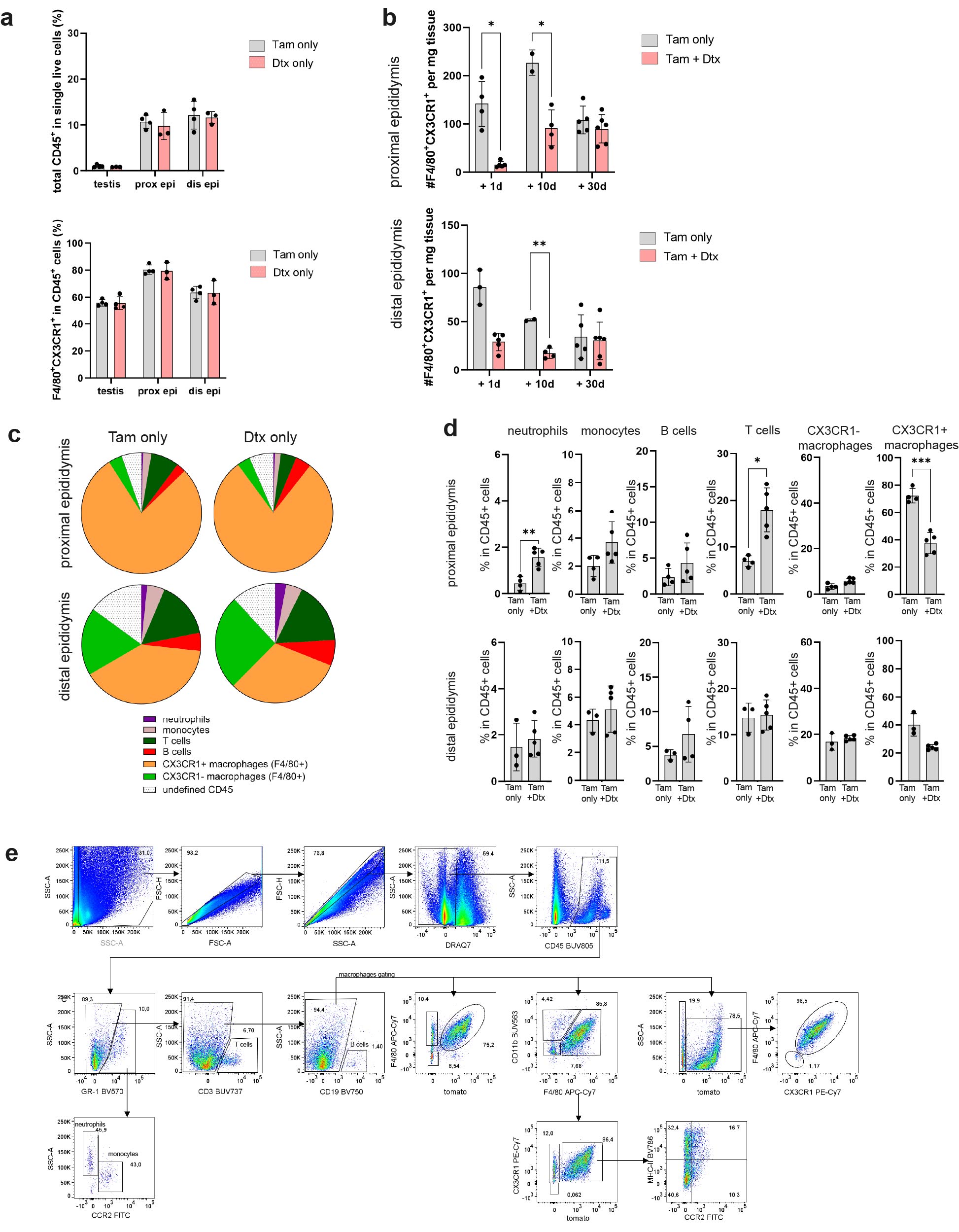

### Supplemental Figure 2

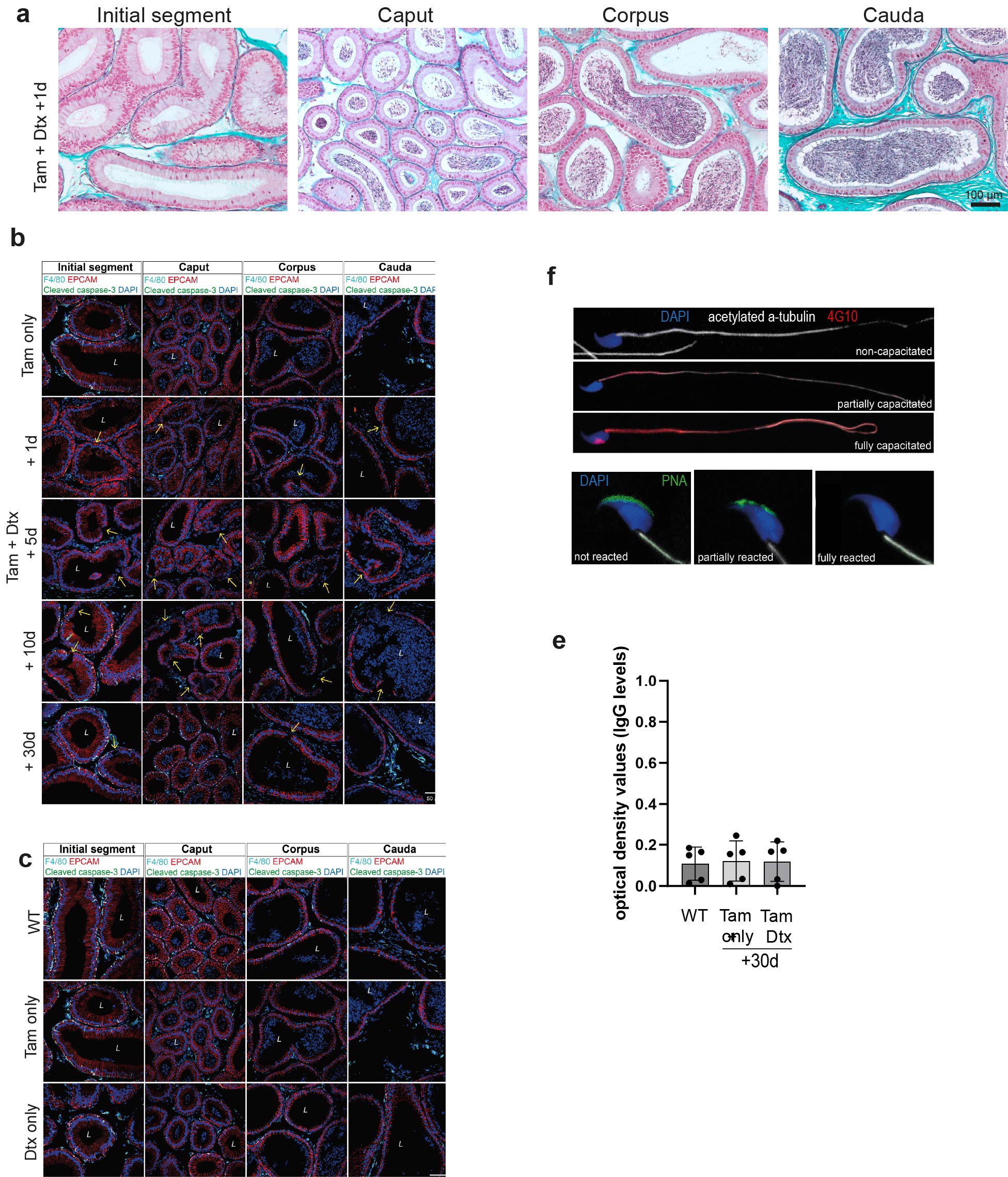

### Supplemental Figure 3

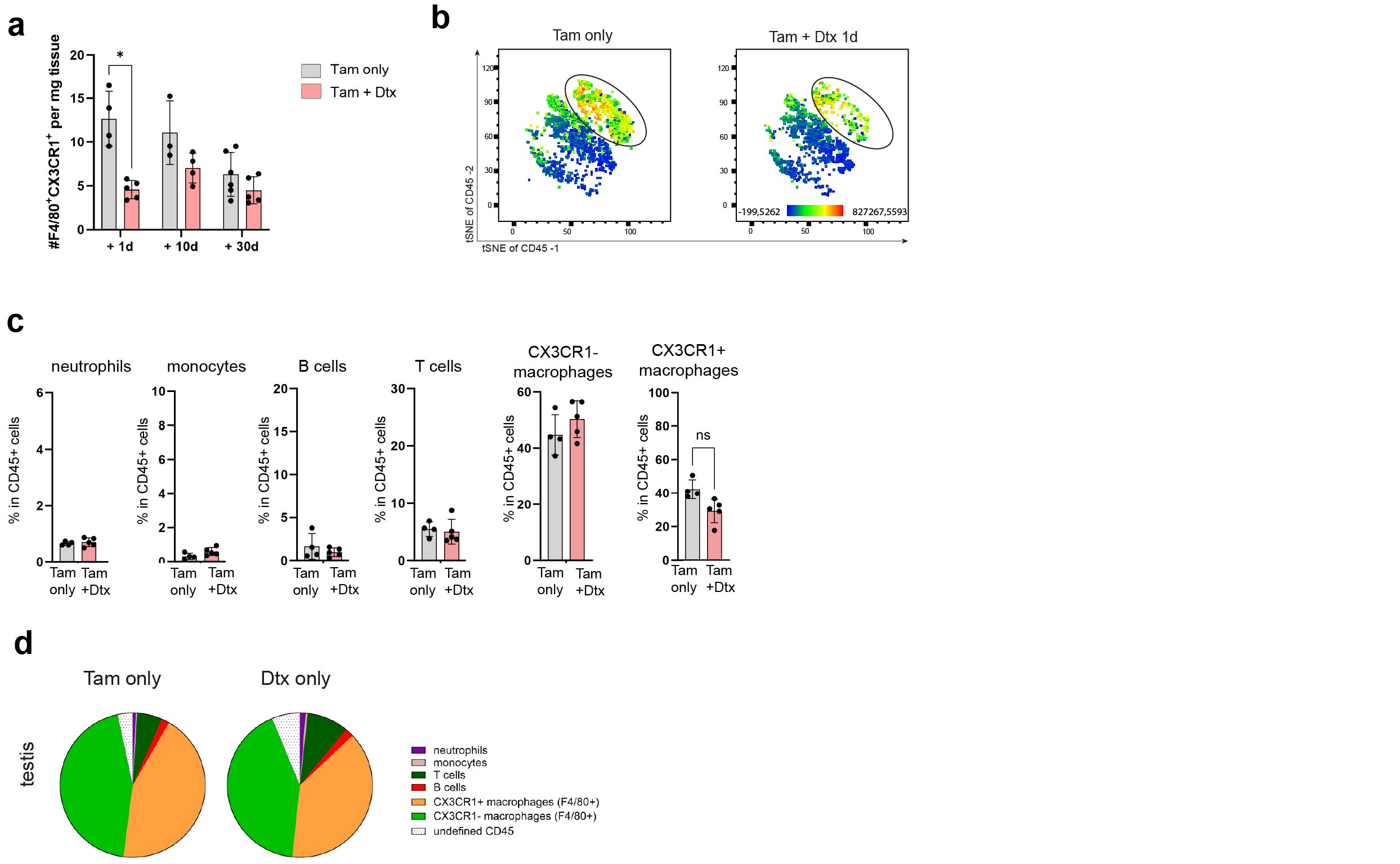

### Supplemental Figure 4

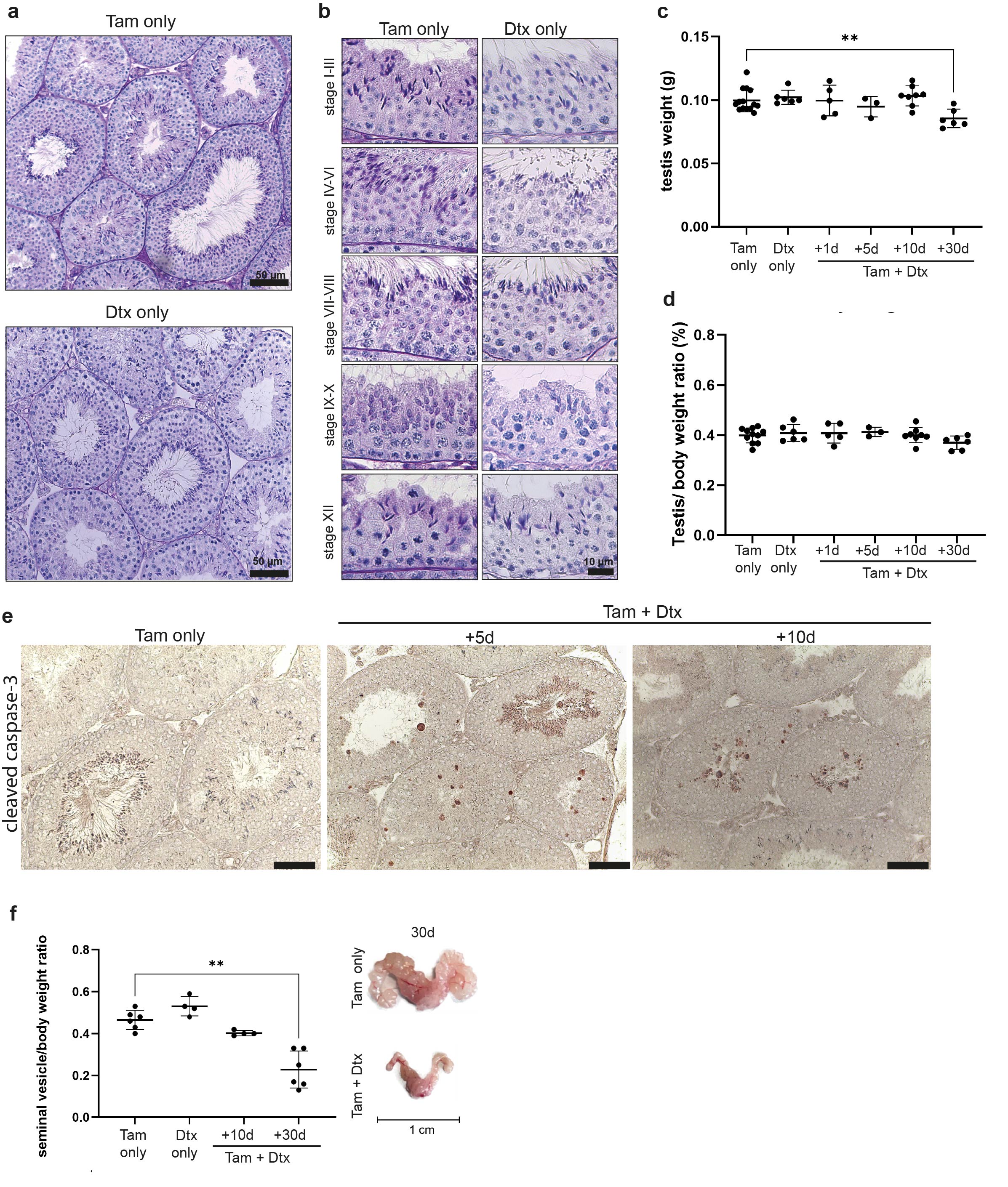

### Supplemental Figure 5

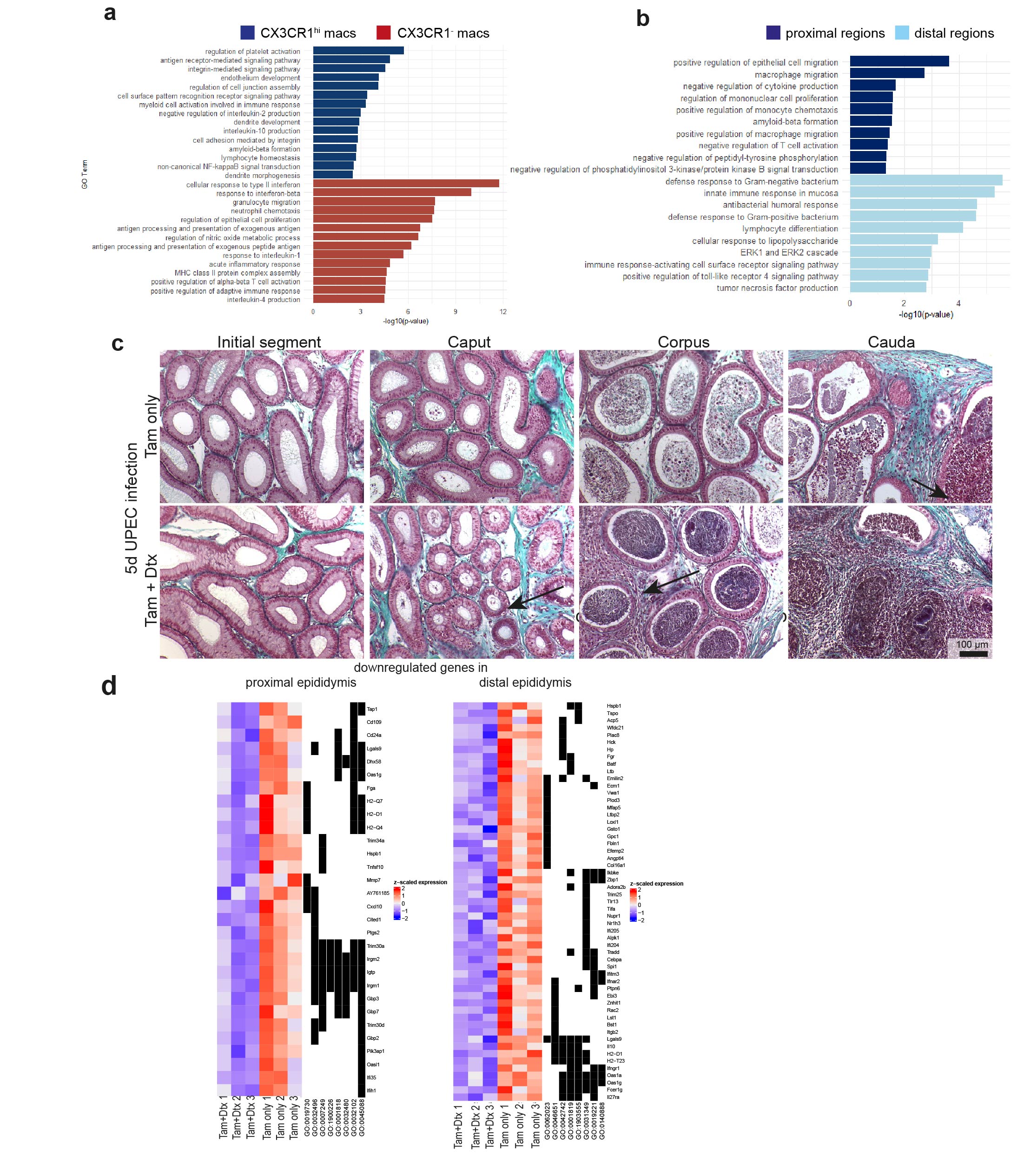

### Supplemental Figure 6

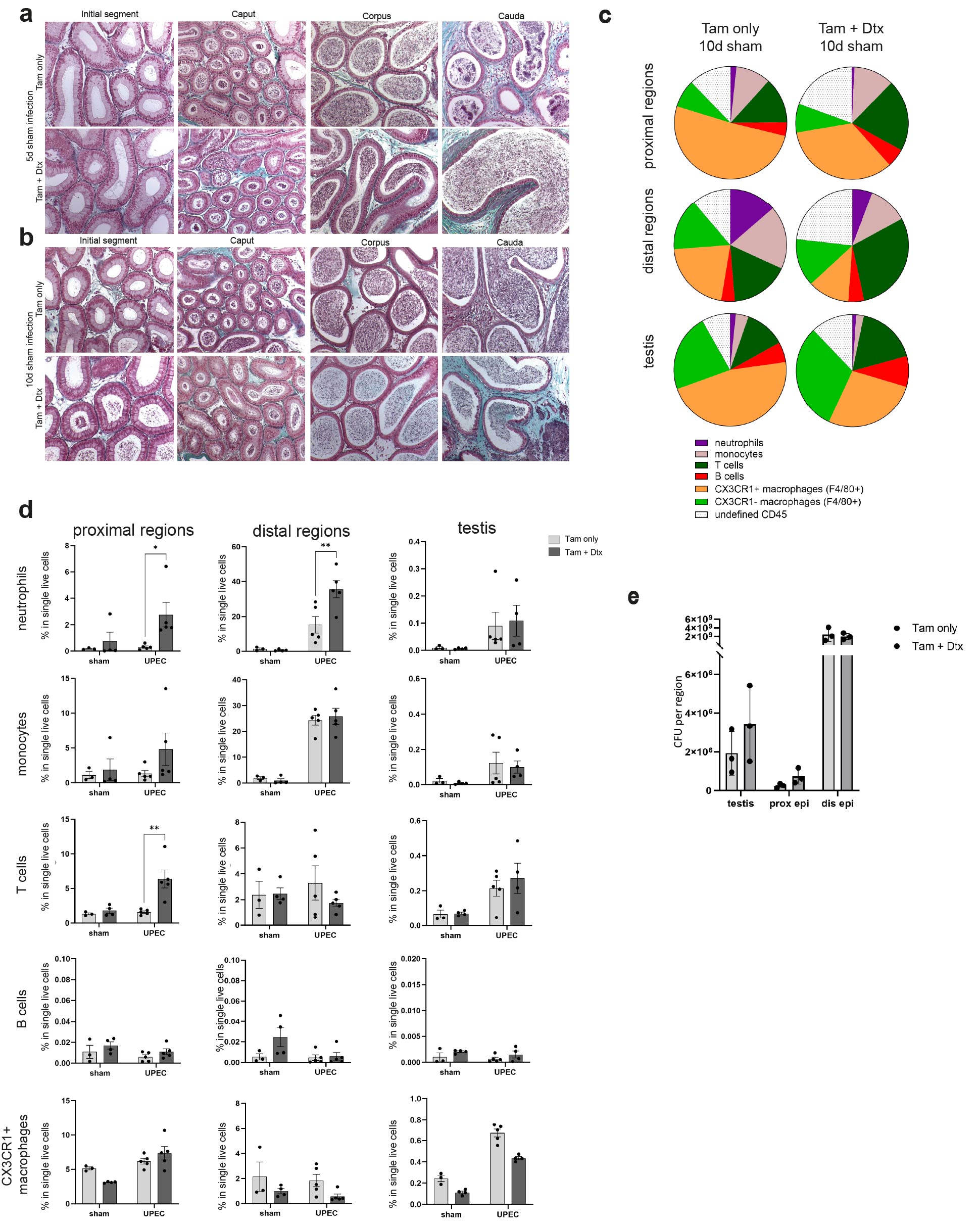

### Supplemental Figure 7

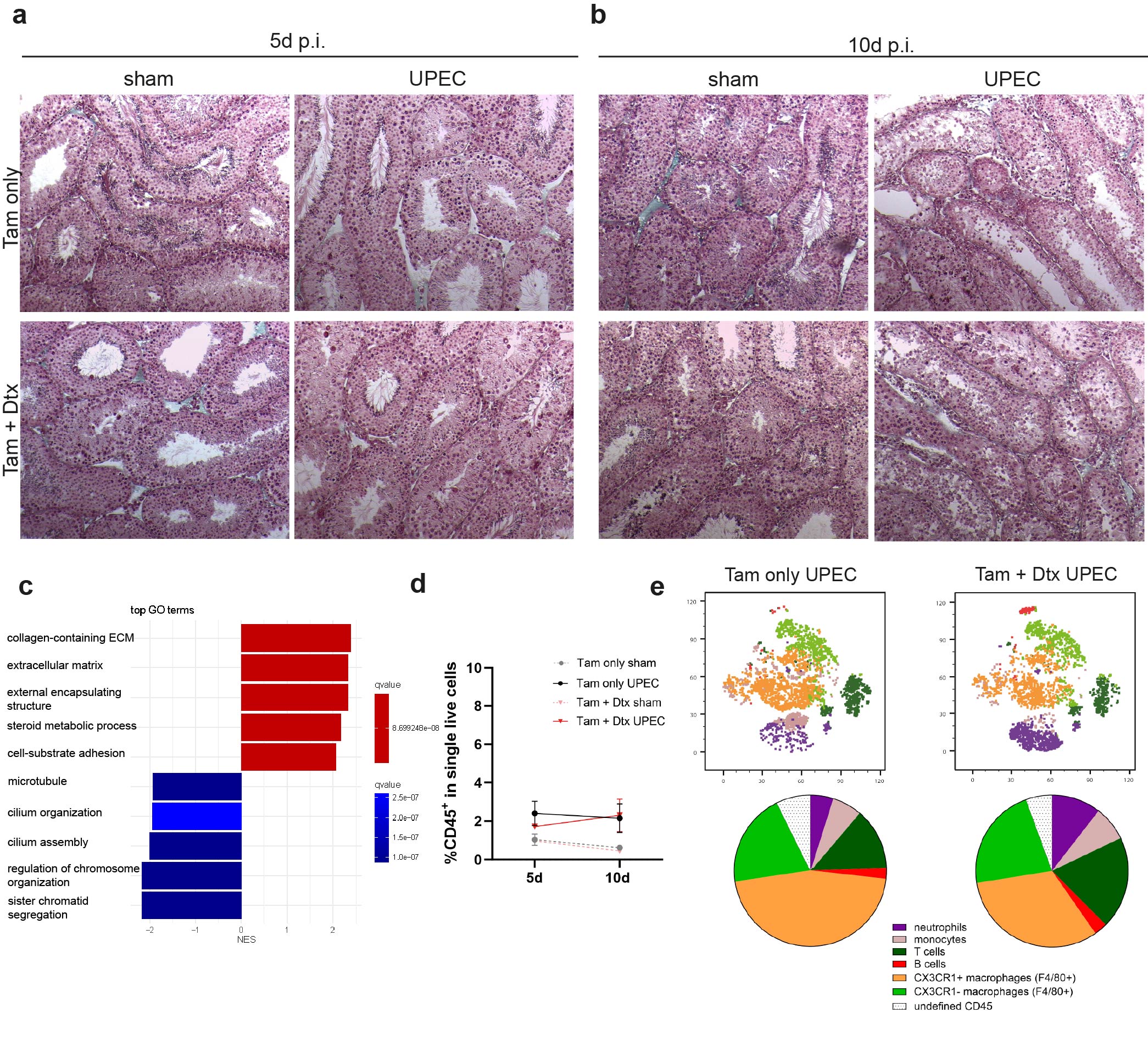
